# Improved transcriptome sampling pinpoints 26 paleopolyploidy events in Caryophyllales, including two paleo-allopolyploidy events

**DOI:** 10.1101/143529

**Authors:** Ya Yang, Michael J. Moore, Samuel F. Brockington, Jessica Mikenas, Julia Olivieri, Joseph F. Walker, Stephen A. Smith

## Abstract

- Studies of the macroevolutionary legacy of paleopolyploidy are limited by an incomplete sampling of these events across the tree of life. To better locate and understand these events, we need comprehensive taxonomic sampling as well as homology inference methods that accurately reconstruct the frequency and location of gene duplications.
- We assembled a dataset of transcriptomes and genomes from 169 species in Caryophyllales, of which 43 were newly generated for this study, representing one of the densest sampled genomic-scale datasets yet available. We carried out phylogenomic analyses using a modified phylome strategy to reconstruct the species tree. We mapped phylogenetic distribution of paleopolyploidy events by both tree-based and distance-based methods, and explicitly tested scenarios for paleo-allopolyploidy.
- We identified twenty-six paleopolyploidy events distributed throughout Caryophyllales, and using novel techniques inferred two to be paleo-allopolyploidy.
- Through dense phylogenomic sampling, we show the propensity of paleo-polyploidy in the clade Caryophyllales. We also provide the first method for utilizing transcriptome data to detect paleo-allopolyploidy, which is important as it may have different macro-evolutionary implications compared to paleo-autopolyploidy.

## Introduction

The prevalence and evolutionary consequences of paleopolyploidy in plants have been discussed at length in the fields of macroevolution (Soltis *et al*., 2015; Lohaus & Van de Peer, 2016). Paleopolyploidy has been correlated with acceleration of speciation (Tank *et al*., 2015; Smith *et al*., 2017), surviving mass extinction (Fawcett *et al*., 2009; Vanneste *et al*., 2014a), evolutionary innovations (Vanneste *et al*., 2014b; Edger *et al*., 2015), and niche shift (Smith *et al*., 2017). While there is little disagreement about the importance of paleopolyploidy in angiosperm evolution, the frequency and phylogenetic locations of these events still remain unclear in many cases. Several limitations in methodology and sampling have limited our ability to accurately locate paleopolyploidy events.

Until recently, most studies of paleopolyploidy have employed either dating synonymous distances (Ks) among paralogous gene pairs (Vanneste *et al*., 2013) or ancestral character reconstruction of chromosome counts (Mayrose *et al*., 2010; Glick & Mayrose, 2014). While these have facilitated the discovery of many paleopolyploidy events, both are indirect methods that have insufficient resolution and can be misleading (Kellogg, 2016). Ks plots between syntenic blocks from individual sequenced genomes have the advantage of being sensitive enough to detect very ancient events (Jaillon *et al*., 2007; Jiao *et al*., 2011; Jiao *et al*., 2012; Jiao *et al*., 2014). However, this technique suffers from the typically sparse taxon sampling available in whole genome data. Distribution of paleopolyploidy events inferred using Ks plots from genomic data, whether or not taking synteny into consideration (Fawcett *et al*., 2009; Vanneste *et al*., 2014a), await re-examination with more comprehensive taxon sampling. An alternative to Ks plots is the detection of polyploidy from chromosome counts. This method has the best signal for recent events and is most often restricted to the genus level or below (Wood *et al*., 2009; Mayrose *et al*., 2010; Mayrose *et al*., 2011).

Recent advance in transcriptome sequencing offers the ability to not only measure Ks distances but also use gene tree topology to validate these. This has allowed for the identification and placement of paleopolyploidy events across the tree of life (Cannon *et al*., 2015; Edger *et al*., 2015; Li *et al*., 2015; Yang *et al*., 2015; Huang *et al*., 2016; Xiang *et al*., 2016). Despite this rapid increase in the number and precision of mapped paleopolyploidy events, the sampling strategy for many of these studies was aimed at resolving deeper phylogenetic relationships. Testing hypotheses regarding the rich macroevolutionary legacy of paleopolyploidy requires much more extensive sampling of genomes and transcriptomes within a major plant clade. To date, very few such data sets with sufficient sampling are available (except Huang *et al*., 2016; Xiang *et al*., 2016). Furthermore, most of these studies have assumed autopolyploidy and have not explicitly tested for allopolyploidy (Kellogg, 2016), primarily because sampling has not allowed for more explicit examination of allopolyploidy. Despite the rich body of literature on gene expression, transposon dynamics, formation of novel phenotypes, and gene silencing and loss in recently formed allopolyploids (reviewed by Soltis & Soltis, 2016; Steige & Slotte, 2016), the long-term effects of paleo-allopolyploidy event remained poorly known (except Estep *et al*., 2014).

The plant order Caryophyllales offers an excellent opportunity to explore phylogenomic processes in plants. Caryophyllales forms a well-supported clade of ~12,500 species distributed among 39 families (Byng *et al*., 2016; Thulin *et al*., 2016), with an estimated crown age of approximately 67-121 Ma (Bell *et al*., 2010; Moore *et al*., 2010; Smith *et al*., 2017). Species of the Caryophyllales are found on every continent including Antarctica and in all terrestrial ecosystems as well as aquatic systems, occupying some of the most extreme environments on earth, including the coldest, hottest, driest, and most saline ecosystems inhabited by vascular plants. Familiar members of the group include cacti, living stones, a diverse array of carnivorous plants (e.g., the sundews, Venus flytrap, and tropical pitcher plants), and several important crop plants (e.g. beet, spinach, amaranth, and quinoa). Such extraordinary diversity makes Caryophyllales a prime system for investigating the relationship between paleopolyploidy and species diversification rate, character evolution, and niche shift. Previous analyses using transcriptomes representing 67 species across Caryophyllales located 13 paleopolyploidy events (Yang *et al*., 2015). By generating 43 new transcriptomes we have expanded the previous sampling to include lineages with key evolutionary transitions, across a dataset that now includes 169 species of Caryophyllales.

As transcriptome datasets grow, all-by-all homology searches become impractical. Hence, in this study, we use a “modified phylome” strategy to build homolog and ortholog groups for species tree inference. In addition, we use an all-by-all approach to build lineage-specific homolog gene sets (Yang & Smith, 2014), and take advantage of recent developments in gene tree-based methods for mapping paleopolyploidy events (Cannon *et al*., 2015; Li *et al*., 2015; Yang *et al*., 2015). Our dense sampling allows us to take chromosome counts into consideration, and begin to explore ancient allopolyploidy events. These improved methods for tree building and mapping of gene duplications, along with our improved taxon sampling, enable the most extensive exploration of paleopolyploidy yet attempted in a major angiosperm clade. The results reported here help establish the necessary foundation for further exploring the macroevolutionary consequences of paleopolyploidy (for example, Smith *et al*., 2017).

## Materials and Methods

### Taxon sampling, laboratory procedure, and sequence processing

We included 178 ingroup datasets (175 transcriptomes, 3 genomes; Table S1) representing 169 species in 27 out of the 39 Caryophyllales families (Byng *et al*., 2016; Thulin *et al*., 2016). Among these, 43 transcriptomes were newly generated for this study (Table S2). In addition, 40 outgroup genomes across angiosperms were used for rooting gene trees (Table S1). Tissue collection, RNA isolation, library preparation, sequencing, assembly, and translation followed previously published protocols (Brockington *et al*., 2015; Yang *et al*., 2017) with minor modifications (Tables S1–S2).

### Caryophyllales homology and orthology inference from peptide sequences using a “modified phylome” strategy

We employed a modified phylome strategy for reconstructing orthogroups. An “orthogroup” includes the complete set of genes in a lineage from a single copy in their common ancestor.

Each node in an orthogroup tree can represent either a speciation event or a gene duplication event. The modified phylome procedure consisted of two major steps. First, “backbone homolog groups” were constructed using peptide sequences from all 43 genomes (3 in Caryophyllales and 40 outgroups). Second, peptides from transcriptomes were sorted to each backbone homolog. This two-step procedure allowed us to avoid the computationally intensive all-by-all homology search for constructing orthogroups.

To construct the backbone homolog groups, we started from the beet proteome (Dohm *et al*., 2014; http://bvseq.molgen.mpg.de/ v1.2, accessed June 25, 2015). Sequences from each beet locus was used to search against a database consisted of combined proteomes from all 43 genomes using SWIPE v2.0.11 (Rognes, 2011) with an E value cutoff of 0.01. The top 100 hits with bitscores higher than 50, and bitscores of at least 20% of the self-hit were retained and aligned using MAFFT v7.215 (Katoh & Standley, 2013), with “--genafpair --maxiterate 1000”. The alignments were trimmed using Phyutility v2.2.6 (Smith & Dunn, 2008) with “-clean 0.1”, and trees were constructed using RAxML v8.1.5 (Stamatakis, 2014) with the model PROTCATWAG. In the resulting trees, terminal branches that were longer than 2 (absolute cutoff) or longer than 1 and more than ten times as long as its sister (relative cutoff) were trimmed; internal branches longer than 1 were separated, leaving only the subtree with the bait beet locus (Yang & Smith, 2014). For each of these trees, a Caryophyllales orthogroup that contained the beet bait locus was recorded. If any of these orthogroups shared beet locus IDs (i.e. had gene duplication within Caryophyllales), the homolog groups were combined, resulting in the final backbone homolog groups.

Next, peptide sequences from each of the 175 Caryophyllales transcriptomes were reduced using CD-HIT v4.6 (-c 0.99-n 5; Fu *et al*., 2012) and sorted to backbone homolog groups using the top SWIPE hit against the beet proteome. A new tree representing each expanded homolog group was estimated using the same settings as for the backbone homolog tree. To reduce isoforms in transcriptome datasets, monophyletic and paraphyletic tips that belonged to the same taxon were removed, leaving only the tip with the highest number of characters in the trimmed alignment (Yang & Smith, 2014). Spurious tips and long internal branches were cut as for the backbone tree. For homolog groups with more than 1,000 and less than 5,000 sequences, alignments were constructed using PASTA v1.6.3 (Mirarab, Siavash *et al*., 2014) with default settings, were trimmed by Phyutility with “-clean 0.01”, and phylogenetic trees were estimated using FastTree v2.1.8 (Price *et al*., 2010) with the model “WAG”. An initial internal branch length cutoff of 2 was used after reducing tips and trimming spurious tips with the same cutoffs as for the backbone trees. A second round of alignment and refining was carried out for these larger homolog groups. Homolog groups larger than 5,000 were ignored.

After modified phylome, we carried out orthology inference following the “rooted ingroup” method in Yang and Smith (2014). Briefly, for each rooted final Caryophyllales orthogroup, we walked from the root towards the tip. When two daughter nodes share one or more taxa, the side with a smaller number of taxa was separated and both clades were taken into account in the next round until all subtrees contained only one sequence per taxon. For each resulting tree with at least 160 taxa, sequences were pooled, re-aligned using PRANK v140110 (Löytynoja & Goldman, 2010) with default settings, trimmed with Phyutility with “-clean 0.3”, and a new ortholog tree estimated using RAxML with “PROTCATAUTO”. A set of more stringent cutoffs was used to produce the final ortholog trees: absolute tips cutoff of 0.6, relative tip cutoff of 0.3, and an internal branch cutoff of 0.4. Aligned sequences were pooled according to remaining tips, trimmed (Phyutility with “-clean 0.3”), and remaining alignments with at least 150 characters and 160 taxa were used for species tree inference.

### All-by-all homology search and orthology inference in each of five Caryophyllales subclades from coding sequences (CDS)

Although modified phylome produced Caryophyllales-wide ortholog trees, uncertainty in alignment and tree inference becomes significant as the dataset size grows. Given the absence of paleopolyploidy events along the backbone of Caryophyllales (Smith *et al*., 2015; Yang *et al*., 2015), we divided Caryophyllales into five subclades according to previous phylogenetic analysis (Yang *et al*., 2015): (1) PHYT: Aizoaceae+the “Phytolaccoid clade” that consists of Nyctaginaceae, Phytolaccaceae s.l. (i.e., including *Agdestis)*, Petiveriaceae, and Sarcobataceae (Yang *et al*., 2015), with *Stegnosperma halimifolium* (Stegnospermataceae) and the three Caryophyllales genomes *Beta vulgaris* (beet, Chenopodiaceae; Dohm *et al*., 2014), *Spinacia oleracea* (spinach, Chenopodiaceae; Dohm *et al*., 2014), and *Dianthus caryophyllus* (carnation, Caryophyllaceae; Yagi *et al*., 2014) as outgroups; (2) PORT: the “Portullugo clade” that consists of Molluginaceae+Portulacineae (Edwards & Ogburn, 2012) with the three Caryophyllales genomes as outgroups; (3) AMAR: Amaranthaceae+Chenopodiaceae, with carnation and *Phaulothamnus spinescens* (Achatocarpaceae) as outgroups; (4) CARY: Caryophyllaceae, with spinach, beet and *Phaulothamnus spinescens* (Achatocarpaceae) as outgroups; and (5) NCORE: the clade that is sister to the rest of Caryophyllales, with all three Caryophyllales genomes plus *Microtea debilis, Physena madagascariensis* and *Simmondsia chinensis* as outgroups.

An all-by-all approach was used for homology inference in each subclade following Yang and Smith (2014) with minor modifications (Methods S1). The final alignments from homolog trees with no taxon duplication (i.e. one-to-one orthologs), no more than 1 missing taxon (except requiring full taxon occupancy for CARY and PHYT), and average bootstrap value of at least 80 were trimmed with Phyutility “-clean 0.5”. Trimmed alignment with at least 300 columns were the final orthologs.

### Species tree inference

We used two alternative approaches for constructing species trees for both the entire Caryophyllales using peptides (“modified phylome dataset”) and each of the five subclades using CDS (“the subclade dataset”). First, a supermatrix was constructed by concatenating trimmed ortholog alignments. A ML tree was estimated from the supermatrix using RAxML, partitioning each locus, with the model set to PROTCATAUTO for peptides and GTRCAT for coding sequences for each individual partition. Node support was evaluated by the Internode Certainty All (ICA) scores (Salichos *et al*., 2014) calculated in RAxML using final ortholog trees as input. Probabilistic correction was used to take incomplete taxon occupancy into consideration (Kobert *et al*., 2016; Stamatakis, 2016). As implementation of ICA score calculation was updated in more recent releases of RAxML, we used RAxML v. 8.2.9 for calculating ICA scores.

In addition to the concatenated analyses, we also searched for the Maximum Quartet Support Species Tree (MQSST) using ASTRAL-II v. 4.10.12 (Mirarab, S. *et al*., 2014; Mirarab & Warnow, 2015) starting from ML trees estimated from individual orthologs. Tree uncertainty was evaluated by using 100 multi-locus bootstrap replicates (Seo *et al*., 2005; Seo, 2008; Mirarab, S. *et al*., 2014), starting from 200 fast bootstrap trees for each final ortholog calculated in RAxML.

### Mapping paleopoloyploidy events based on subclade orthogroup tree topology

To map paleopolyploidy events in each subclade, we extracted orthogroups from each subclade homolog tree, requiring no more than two missing ingroup taxa and an average bootstrap percentage of at least 50. For each node in these filtered subclade orthogroup trees, when two or more taxa overlapped between the two daughter clades, a gene duplication event was recorded to the most recent common ancestor (MRCA) on the subclade species tree (Yang *et al*., 2015). In this procedure, each node on a species tree can be counted at most once per orthogroup to avoid nested gene duplication inflating the number of duplication scored. In addition to applying a bootstrap filter, we also used a second, local topology filter (Cannon *et al*., 2015; Li *et al*., 2015) that only mapped a gene duplication event when the sister clade of the gene duplication node in the orthogroup contained a subset (or all) of the taxa in the corresponding sister clade in the subclade species tree.

### Distribution of synonymous distance gene pairs (Ks plots)

For each of the ingroup Caryophyllales datasets, a Ks plot of within-taxon paralog pairs was created following the same procedure as Yang et al. (2015) based on BLASTP hits. Similarly, we carried out a second Ks analysis based on BLASTN between CDS without first reducing highly similar sequences to maximize detection of more recent paleopolyploidy events. In addition to within-taxon Ks plots, we also calculated Ks distribution of between-species reciprocal best BLASTN hit pairs.

### Chromosome counts

Chromosome counts were obtained from the Chromosome Counts Database (http://ccdb.tau.ac.il/Angiosperms/Caryophyllaceae/ accessed Oct 5, 2015). When counts in this database were unavailable or highly inconsistent, counts were obtained from the Jepson eFlora (http://ucjeps.berkeley.edu/eflora/ accessed Oct 5, 2015) and Flora of North America (http://www.efloras.org/ accessed Oct 5, 2015).

### Total evidence approach for mapping paleopolyploidy events

We consider six scenarios for mapping paleopolyploidy events taking orthogroup tree topology, Ks plots, and chromosome counts into consideration (Fig. 1). When paleopolyploidy events occurred without subsequent speciation (or for which only one taxon is represented in our sampling), because we masked tips to accommodate isoforms from transcriptomes and because we required at least two overlapping taxa between sister clades to record a gene duplication event, these paleopolyploidy events could only be detected from a Ks peak unique to a terminal branch (Fig. 1, a-c; Kellogg, 2016). When two or more taxa diversified following an paleopolyploidy event, it could be detected from both a shared Ks peak and elevated gene duplications in their MRCA (Fig.1d; Yang *et al*., 2015). When not all taxa in the clade mapped by gene duplications shared the Ks peak, it could due to either allopolyploidy or phylogenetic uncertainty (Fig. 1, e and f). We did not consider polyploidy events that were supported by a single chromosome count alone, as these can be more recent genome duplication events, chromosome number can vary even within population (Caperta *et al*., 2016), and an increase in number can represent chromosome fission instead of duplication (Fishman *et al*., 2014; Chester *et al*., 2015).

**Figure 1.**
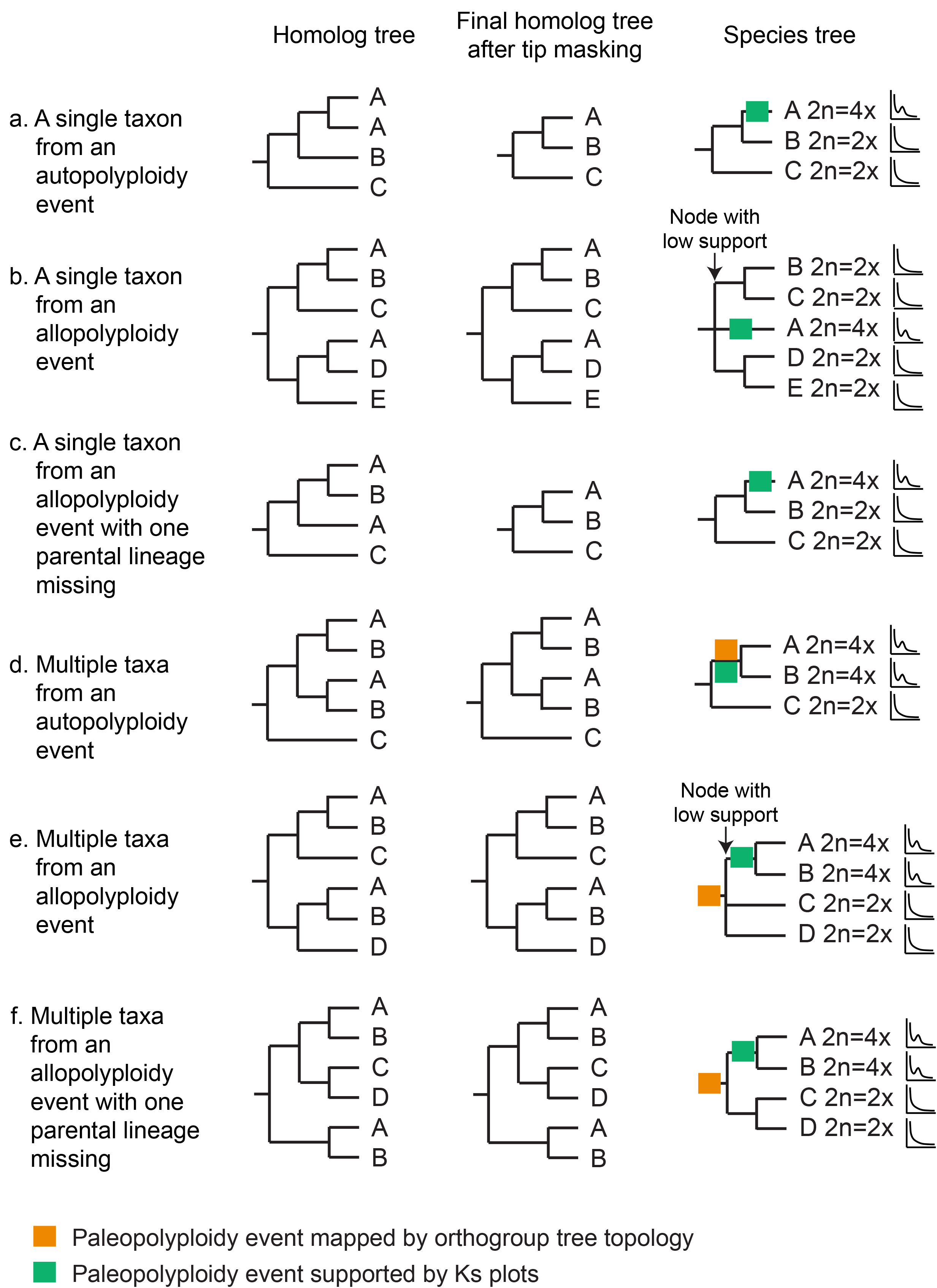
Scenarios of paleopolyploidy events. Letters A-E represent taxon names, followed by chromosome numbers with the base number X, and schematic Ks plots (x axis are Ks values, and y axis are number of paralogous gene pairs). Modified from Kellogg (2016)

## Results

### Data availability

Raw reads for newly generated transcriptomes were deposited in the NCBI Sequence Read Archive (BioProject: PRJNA388222; supplementary table S2). Assembled sequences, alignments, and trees were deposited in Dryad (doi:XXXX). Scripts used were also archived in Dryad, with notes and updates for modified phylome available from https://bitbucket.org/yangya/genome walking 2016 and those for building lineage-specific homologs and mapping paleopolyploidy events available from https://bitbucket.org/blackrim/clustering.

### RAxML and ASTRAL recovered nearly identical species tree topologies

Both analyses using RAxML and ASTRAL recovered identical topologies for most branches (Figs. 2, S1–S3). We consider branches with an ICA score higher than 0.5 as strongly supported, as ICA scores lower than 0.5 suggests that the dominant bipartition is present in less than 80% of ortholog trees (Salichos *et al*., 2014). As multi-locus bootstrap support percentages increase with the number of loci (Seo, 2008) and given that each of our final ortholog set contained more than a hundred loci (Table 1), we consider multi-locus bootstrap values less than 100 as low support. Using this set of criteria, most branches from subclade datasets (Figs. 2 & S1) and the majority of the branches from modified phylome (Figs. S2 & S3) were well-supported.

**Table 1.** Statistics for homology and orthology inference. PHYT, PORT, AMAR, CARY, and NCORE are subclades within Caryophyllales (see Fig. 3). Orthology inference methods are from Yang et al. (2015).

|  | Caryophyllales | PHYT | PORT | AMAR | CARY | NCORE |
| --- | --- | --- | --- | --- | --- | --- |
| Data type | Peptides | Coding sequences (CDS) |  |  |  |  |
| Homology inference | Modified phylome | All-by-all |  |  |  |  |
| Orthology inference | “Rooted ingroup” | One-to-one orthologs |  |  |  |  |
| No. taxa (ingroup+outgroup) | 175+40 | 45+4 | 29+3 | 37+2 | 31+3 | 31+6 |
| Min. taxa per Ortholog | 160 | 49 | 31 | 38 | 34 | 36 |
| Supermatrix dimension taxa x loci (columns) | 178 x 624 (215,669) | 49 x 152 (217,033) | 32 x 171 (230,873) | 39 x 315 (453,842) | 34 x 736 (1,130,082) | 37 x 213 (325,966) |
| Supermatrix gene / character occupancy | 92.6% / 80.1% | 100% / 92.5% | 97.8% / 86.5% | 98.1% / 87.2% | 100% / 92.2% | 97.5% / 87.8% |
| Min. ingroup taxa for mapping gene duplication | N/A | 43 | 27 | 35 | 29 | 29 |
| No. of ingroup clades used for mapping gene duplications (average bootstrap $\geq 50$) | N/A | 2,818 | 3,561 | 4,713 | 6,657 | 1,310 |

**Figure 2.**
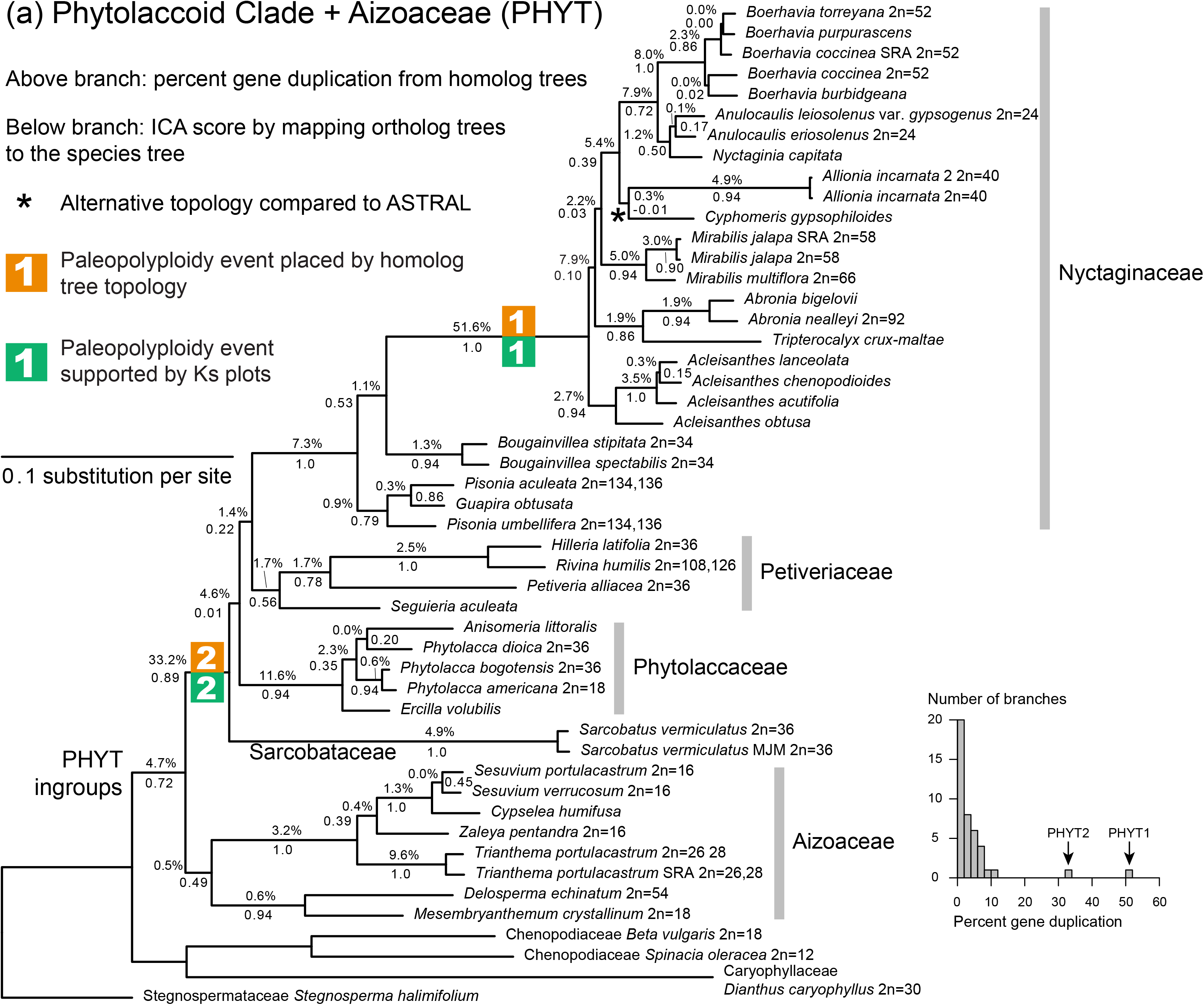
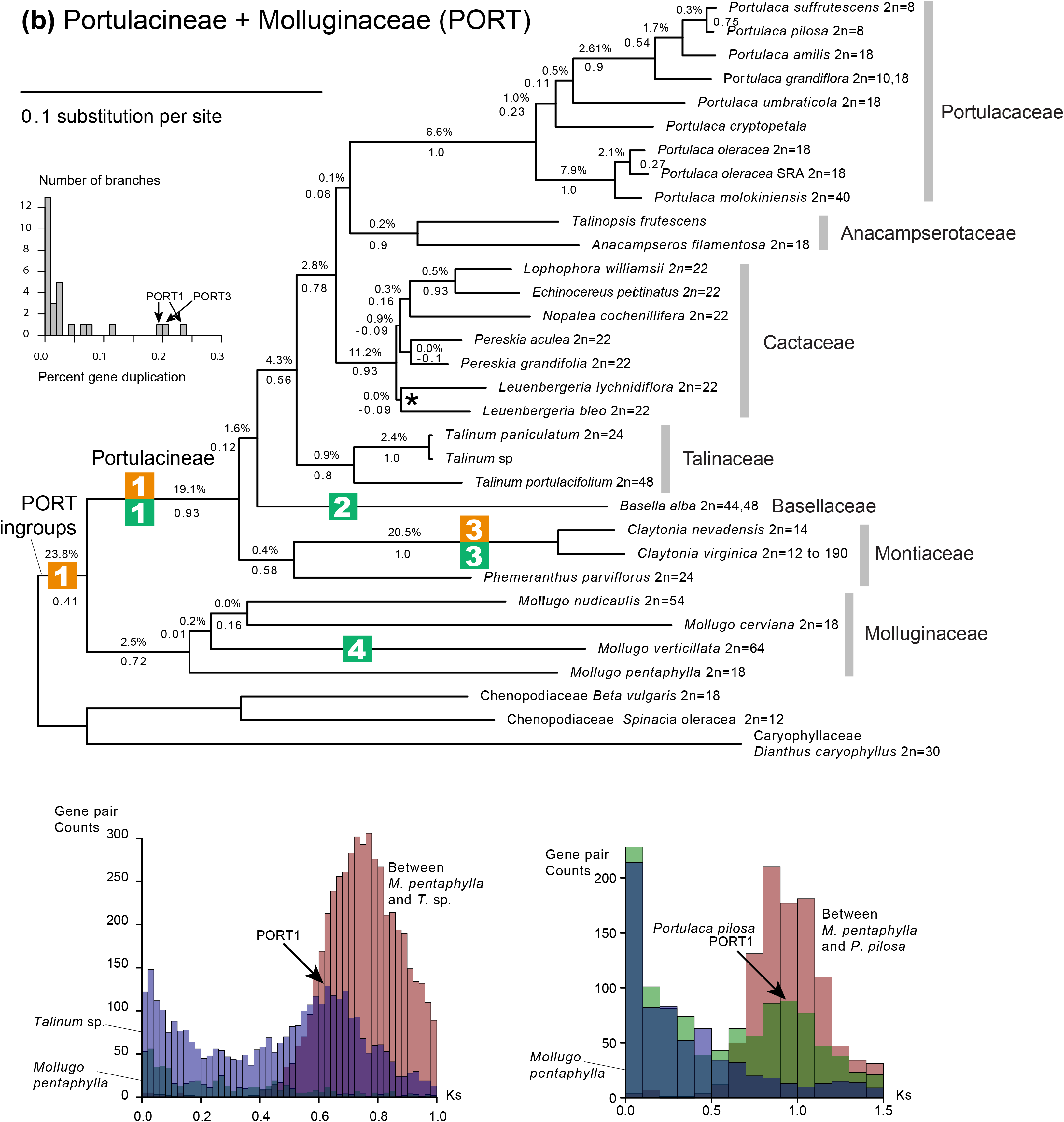
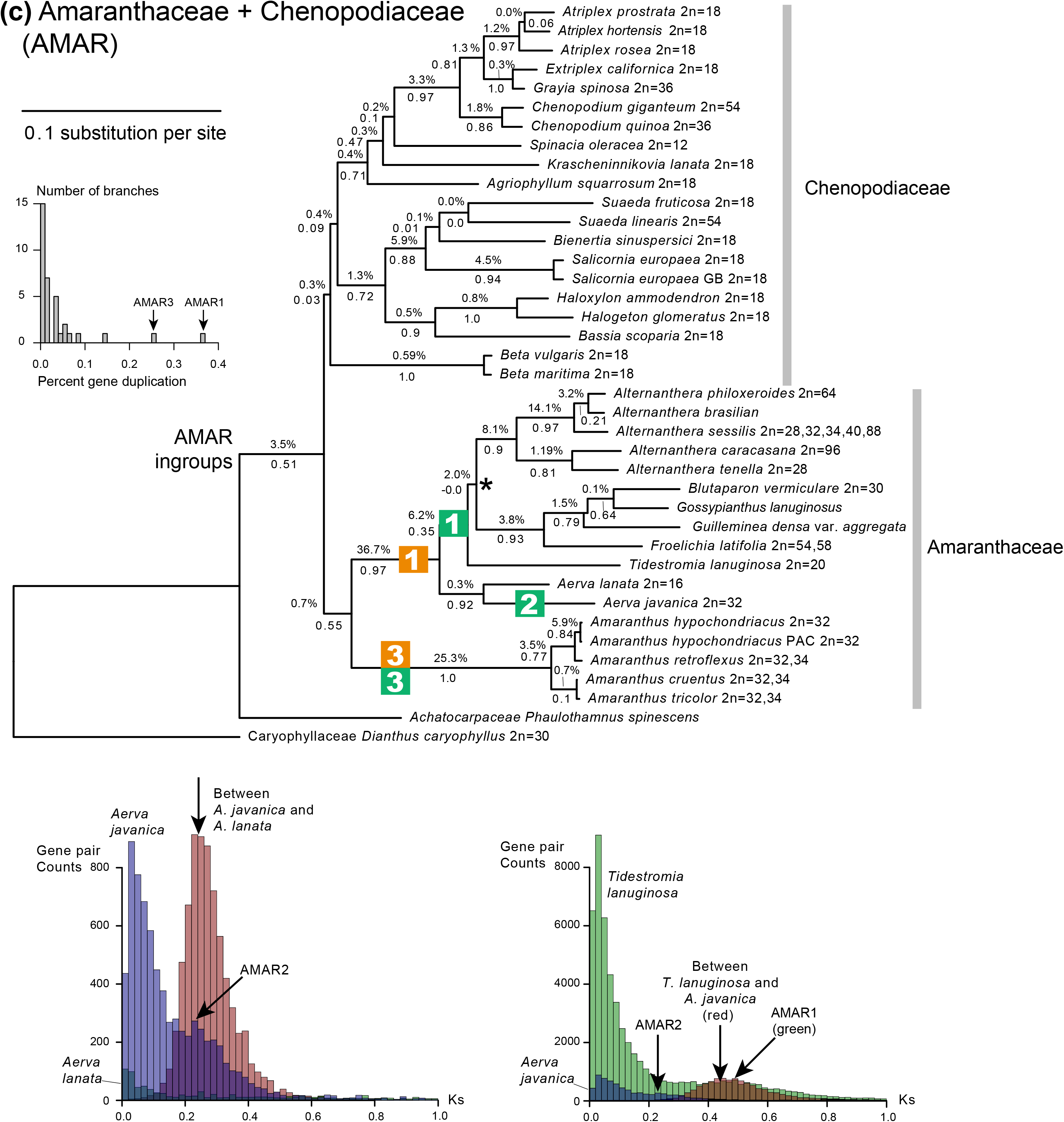
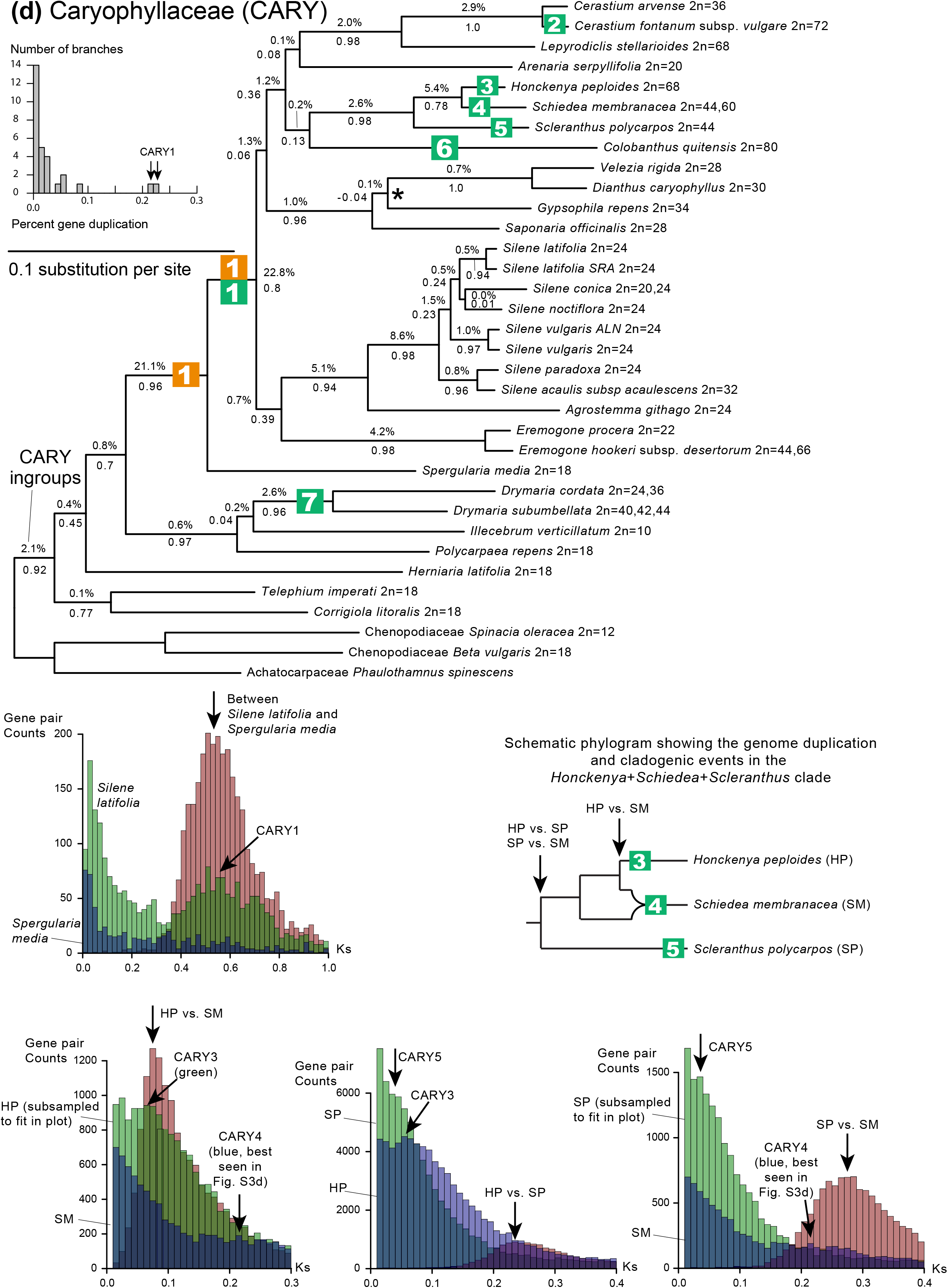
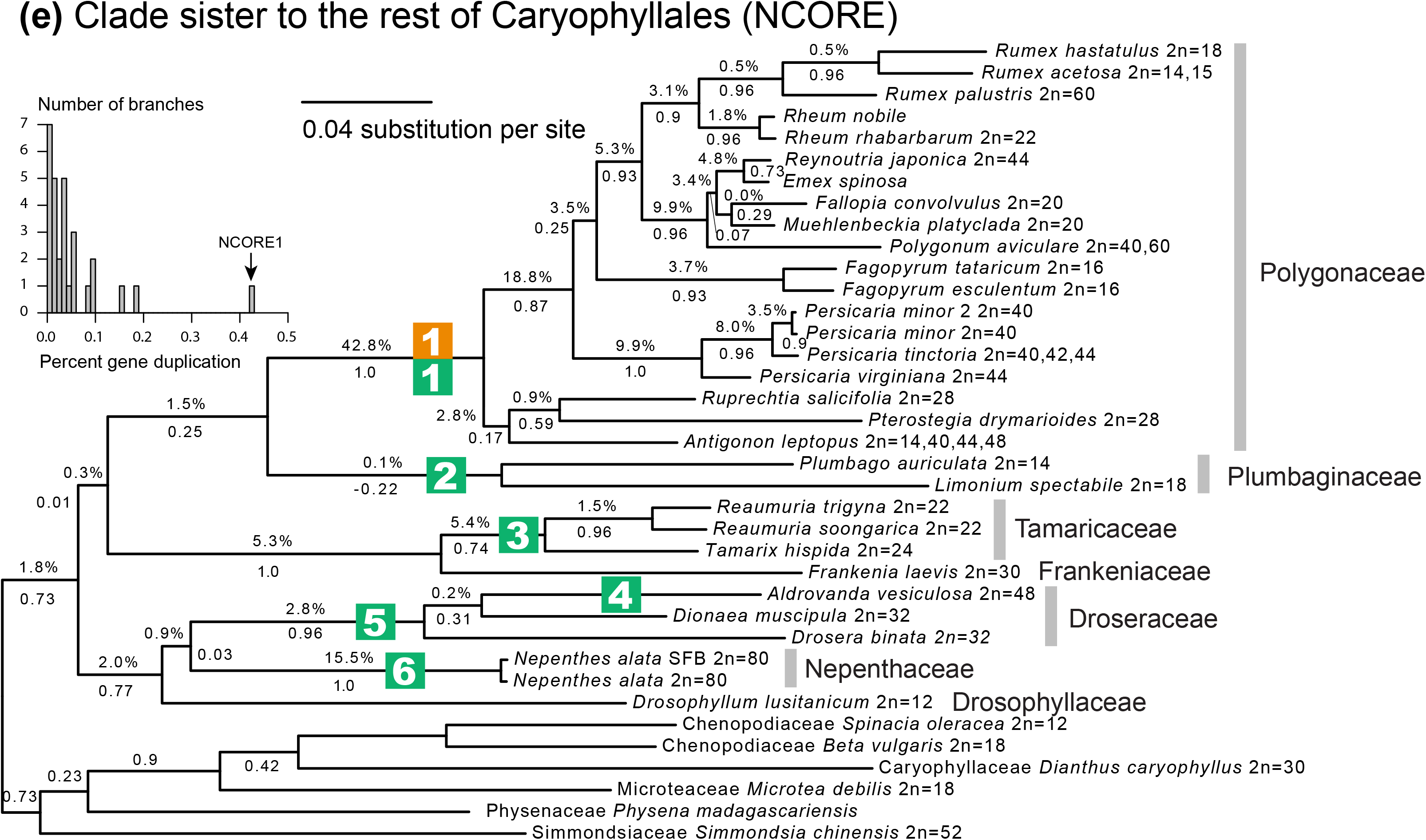
Best tree from RAxML analysis of concatenated supermatrices of subclades. Percentage values on branches indicate proportion of orthogroups showing duplication without filtering by local tree topology. ICA values are given below branches. Selected Ks plots based on BLASTN hits are shown below trees. Ks values lower than 0.01 are not shown. (a) PHYT: the phytolaccoid clade and Aizoaceae; (b) PORT: Portulacineae and Molluginaceae; (c) AMAR: Amaranthaceae and Chenopodiaceae; (d) CARY: Caryophyllaceae; and (e) NCORE: the clade sister to rest of the Caryophyllales.

We recovered between 152 to 736 final one-to-one orthologs and 0.2-1.1 million trimmed CDS columns from each of the five subclades. The concatenated supermatrices had gene occupancies of 98-100% and character occupancies of 87-93% (Table 1). Four clades showed different relationships between RAxML and ASTRAL, with little support for either alternative relationships (Fig. 2 marked with “*”; Figure S1): (1) *Cyphomeris gypsophiloides* (Nyctaginaceae, PHYT) was sister to *Allionia* in the RAxML tree (ICA = -0.01) but was sister to the clade *Nyctaginia+Anulocaulis+Boerhavia* in the ASTRAL tree (bootstrap = 95); (2) species in *Leuenbergeria* (Cactaceae, PORT) were monophyletic in the RAxML tree (ICA = -0.09) but were paraphyletic to the rest of Cactaceae in the ASTRAL tree (bootstrap = 69); (3) *Tidestromia lanuginosa* (Amaranthaceae, AMAR) was sister to the clade of *Froelichia+Guilleminea+Gossypianthus+Blutaparon+Alternanthera* in the RAxML tree (ICA = -0.00), but was sister to *Alternanthera* in the ASTRAL tree (bootstrap = 50); and (4) *Saponaria officinalis* (Caryophyllaceae, CARY) was sister to *Gypsophila+Dianthus+Velezia* in the RAxML tree (ICA = -0.04), but was sister to *Dianthus+Velezia* in the ASTRAL tree (bootstrap = 63).

Among the 15,045 homolog groups we obtained by modified phylome, 15 had more than 5,000 sequences and were ignored, while the rest were used for subsequent orthology inference. The final concatenated matrix consisted of 624 loci and 215,669 amino acids, with a final gene occupancy of 92.6% and character occupancy of 80.1% (Table 1). The modified phylome dataset recovered identical species tree topologies except for one branch that had little support from either analysis (Figs. S2 & S3): *Leuenbergeria* (Cactaceae) was monophyletic in the RAxML tree (ICA = 0.18) but was polyphyletic in the ASTRAL tree (bootstrap = 28). The modified phylome analysis recovered an identical species tree topology compared to that recovered by the subclade analysis. When subclade trees had different topologies between RAxML and ASTRAL, the modified phylome tree agreed with the subclade ASTRAL results in the placement of *Cyphomeris gypsophiloides* (ICA = 0.23 and bootstrap = 97), whereas the position of *Leuenbergeria* was recovered in the same incongruent positions as recovered by RAxML and ASTRAL in the subclade tree analyses. The modified phylome tree recovered both *Tidestromia lanuginosa* (0.19/81) and *Saponaria officinalis* (0.38/98) in the same positions as found in the RAxML results for the subclade trees.

### Twenty-six paleopolyploidy events were mapped

A total of 26 paleopolyploidy events were inferred by using a total evidence approach of orthogroup tree topology, shared Ks peaks, and chromosome counts (Figs. 2 & 3). Overall the two orthogroup tree filtering strategies, with or without considering local tree topology, produced almost identical results for both gene and genome duplications (Figs. 2 & S4). The frequency of gene duplications was strongly associated with the inferred paleopolyploidy events (Fig. 2). In our Ks analysis, we only considered Ks peaks that were similar in height or taller than the peak, around Ks = 2, that corresponds to the early eudicot paleohexaploidy that predates the origin of Caryophyllales (Dohm *et al*., 2012; Jiao *et al*., 2012; Yang *et al*., 2015).

**Figure 3.**
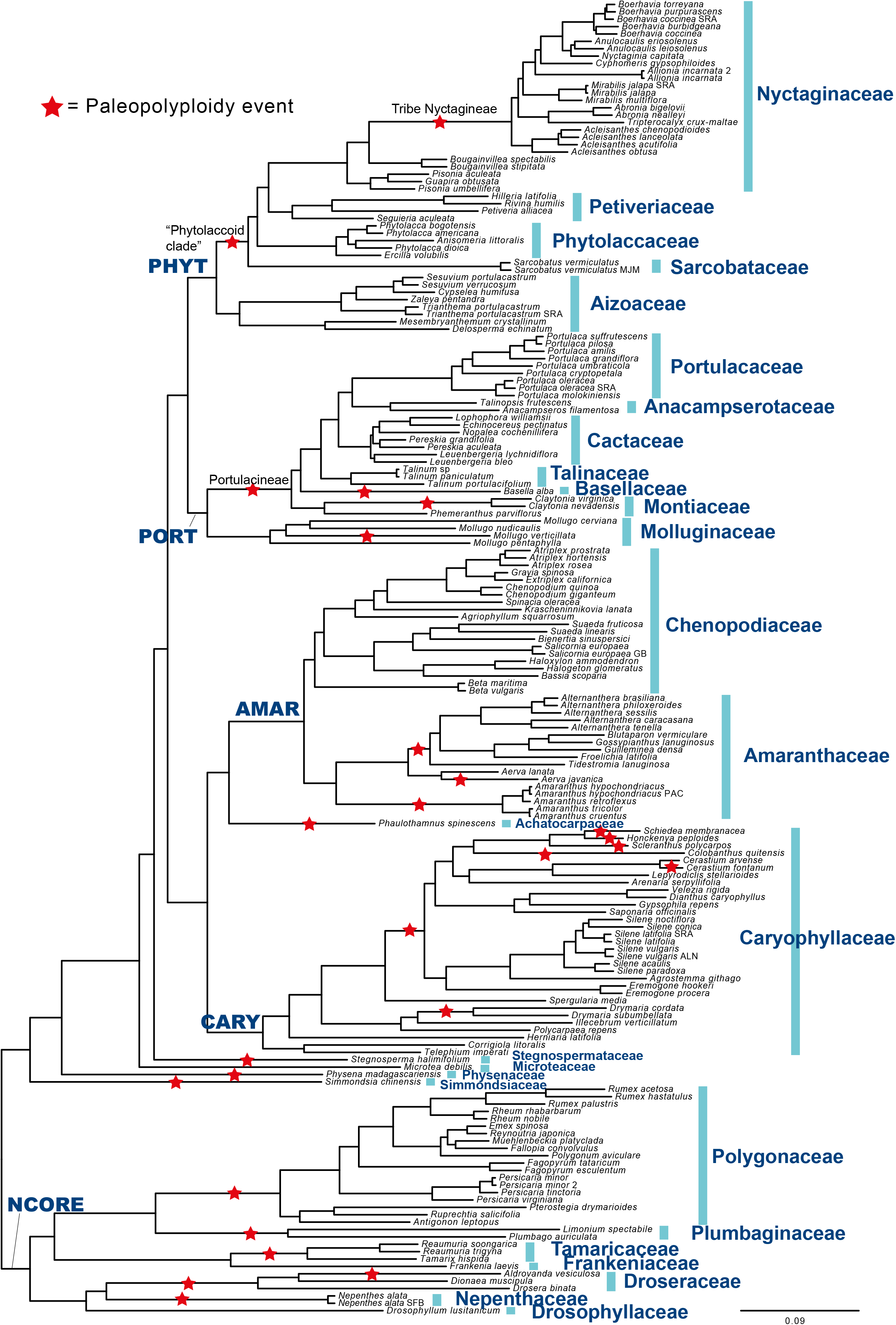
Summary of paleopolyploidy events mapped to the species tree from RAxML analysis of the concatenated 624-gene peptide supermatrix from modified phylome.

Two paleopolyploidy events in the PHYT clade were well supported by both homolog tree topology and shared Ks peaks (Figs. 2a, S4a, and S5a). The percentage of genes with duplications were 52% without and 45% with local filtering for PHYT1, and 33% and 31% respectively for PHYT2. Chromosome counts were relatively high in Nyctaginaceae as compared to the rest of the PHYT clade, but were difficult to use as evidence for paleopolyploidy because they varied among closely related species and were not always multiples of the same base number.

At least four paleopolyploidy events were recovered in the PORT clade. PORT1 was mapped to both the MRCA of Portulacineae (19%/17%) and its parent node (24%/22%) from gene duplications (Figs. 2b and S4b). However, Molluginaceae did not share the Ks peak that was present in all members of Portulacineae (Fig. S5b), and chromosome counts were uninformative for the placement of PORT1. Within-species paralogs in Portulacineae (represented by the PORT1 Ks peak in *Talinum* sp.; Fig. 2b, lower left) coalesced at lower Ks values compared to the Ks peak between *Talinum* sp. and *Mollugo pentaphylla* (a member of Molluginaceae). However, similar comparison of *Portulaca pilosa* (a member of Portulacineae) vs. *Mollugo pentaphylla* showed overlapping Ks peaks (Fig. 2b, lower right), likely due to faster molecular rate in *Portulaca* compared to *Talinum*. Therefore, phylogenetic uncertainty likely at least partly contributed to the ambiguity in mapping and we did not see evidence supporting an allopolyploidy event at the base of Portulacineae. Both PORT2 and 4 were recovered by taxon-specific Ks peaks, and both had relatively high chromosome counts compared to close relatives (Figs. 2b, S4b, and S5b). PORT3 was supported by shared Ks peaks and gene duplications in orthogroup trees (21%/18%), whereas chromosome counts were uninformative. Aside from these four paleopolyploidy events, we also observed slightly elevated gene duplication percentages at the MRCA of Cactaceae (11.2%/6.7%). However, neither Ks plots nor chromosome counts supported a paleopolyploidy event at that node.

At least three paleopolyploidy events were recovered in the AMAR clade (Figs 2c, S4c, and S5c). AMAR1 was detected by an elevated percentage of gene duplications mapped to the branch uniting *Alternanthera+Gossypianthus+Blutaparon+Froelichia+Aerva* (37%/35%). However, species of *Aerva* lacked the Ks peak shared by *Alternanthera+Gossypianthus+Blutaparon+Froelichia* at 0.4-0.65. Further examination of between-species Ks peaks showed that the Ks peak around 0.23 in *Aerva javanica* was more recent than the Ks peak between *Aerva javanica* and *Tidestromia lanuginosa* (approximately 0.6), suggesting that paralogs in *Aerva javanica* coalesced within *Aerva* before coalescing with taxa outside of *Aerva*, and AMAR1 and 2 were two distinctive genome duplication events (Fig. 2c). In addition, the Ks peak within *Aerva javanica* overlapped with the Ks peak of *Aerva javanica* vs. A.*lanata*. Given that *Aerva javanica* had faster molecular substitution rate than *A. lanata* according to their relative branch lengths (Fig. 2c), paralogous copies within *A. javanica* most likely coalesced along the branch leading to *A. javanica* before coalescing with *A. lanata*. The lack of the AMAR2 peak in *A. lanata* as well as chromosome counts (2n = 32 for *A. javanica* vs. 2n = 16 for *A. lanata)* further supported the location of AMAR2 along the terminal branch leading to *A. javanica*. Based on the lack of the AMAR1 peak in both *Aerva* species, we inferred that AMAR1 was an allopolyploidy event, with one parental lineage closely related to *Aerva* and the other parental lineage missing, consistent with the scenario in Fig. 1f.

At least seven paleopolyploidy events were recovered in the CARY clade (Figs. 2d, S4d, and S5d). CARY1 was detected through an elevated percentage of gene duplication in two adjacent nodes (21%/20% on the node that included *Spergularia media*, and 23%/20% in the node excluded *S. media;* Figs. 2d and S4d). However, *S. media* did not share the CARY1 Ks peak. The reciprocal best hits Ks peak between *Spergularia media* and *Silene latifolia* indicated that paralogs derived from CARY1 coalesced within *Silene latifolia* at similar Ks values compared to coalescing with *Spergularia media* (Fig. 2d), suggesting that phylogenetic uncertainty at least partly contributed to the fact that CARY1 mapped to two adjacent nodes.

Following CARY1, five taxa showed more recent Ks peaks nested within CARY1 (Figs. 2d, S4d, and S5d). Among them, CARY2 was observed in *Cerastium fontanum* (2n = 72) but missing from its sister *C. arvense* (2n = 36), and therefore mapped to the terminal branch leading to *C. fontanum. Honckenya peploides* had a Ks peak around 0.06, its sister *Schiedea membranacea* had a Ks peak around 0.2, whereas their reciprocal best hit Ks peak was around 0.08, suggesting that a paleopolyploidy event CARY3 was restricted to the terminal branch leading to *H. peploides*. The observation that the Ks peak within *Schiedea membranacea* (CARY4) was much older than its split with *H. peploides*, but that CARY4 was not shared with *H. peploides*, suggested an allopolyploid origin of *S. membranacea*. Pairwise comparison among *Schiedea membranacea*, *Honckenya peploides*, and *Scleranthus polycarpos* showed that all three each had a paleopolyploidy event restricted to respective terminal branches, and *Schiedea membranacea* likely formed through allopolyploidy with one parental lineage missing from our taxon sampling (Fig. 2d). All three species had relatively high chromosome counts compared to their close relatives.

CARY6, also nested in CARY1, was mapped to the terminal branch leading to *Colobanthus quitensis* that had a chromosome count of 2n = 80, one of the highest in the family. Although *Colobanthus* was weakly supported to be sister to the clade consisted of *Honckenya+Schiedea+Scleranthus* (ICA = 0.13, bootstrap = 100), the peak at approximately Ks = 0.15 was more recent than the *Honckenya/Scleranthus* (Ks~0.24) or the *Schiedea/Scleranthus* (Ks~0.27) split and is therefore inferred to be independent of CARY 3, 4, or 5. A seventh paleopolyploidy event independent of CARY1 was mapped to the MRCA of *Drymaria cordata* and *D. subumbellata* by a shared Ks peak. Both species had relatively high chromosome counts compared to their close relatives.

At least six paleopolyploidy events were inferred in the NCORE clade (Figs. 2e, S4e, and S5e). Among these, both gene duplications (43%/34%) and shared Ks peaks supported a paleopolyploidy event at the base of Polygonaceae (NCORE1). NCORE2-5 were supported by Ks peaks, and NCORE 6 was supported by both Ks peaks and chromosome counts. We inferred NCORE 5 (base of Droseraceae) and NCORE 6 (branch leading to *Nepenthes alata*, Nepenthaceae) as two separate paleopolyploidy events on sister branches given that very low frequencies of gene duplication was mapped to the MRCA of Droseraceae+Nepenthaceae (0.9%/1.7%), compared to the MRCA of Droseraceae (2.8%/3.0%) and Nepenthaceae (16%/15%)

In addition to the paleopolyploidy events detected from each of the five subclades, three of the four taxa along the grade paraphyletic to PHYT+PORT+AMAR+CARY also each had a peak at Ks lower than 1: *Stegnosperma halimifolium* (Fig. S5a), *Physena madagascariensis* (Fig. S5e), and *Simmondsia chinensis* (Fig. S5d). As there is no polyploidy event along the Caryophyllales backbone leading to beet (Chenopodiaceae) from genome analysis (Dohm *et al*., 2012; Dohm *et al*., 2014), any Ks peak mapped to this grade likely represents a lineage-specific paleopolyploidy event. Also, the relatively high chromosome count of *Simmondsia chinensis* (2n = 52) compared to *Microtea debilis* (2n = 18), the only taxon in this grade that did not experience a paleopolyploidy event, further supports the lineage specific nature of the paleopolyploidy events along this grade. In addition, *Phaulothamnus spinescens* (Achatocarpaceae; sister to AMAR) also likely had its lineage specific paleopolyploidy event, as it had a Ks peak that was not shared with its sister clade (Figs. 2c & S5c).

## Discussion

Our analyses add to a growing body of literature that suggests that paleopolyploidy events are much more prevalent than previously thought (Cannon *et al*., 2015; Edger *et al*., 2015; Li *et al*., 2015; Yang *et al*., 2015; Huang *et al*., 2016; Xiang *et al*., 2016; Mandáková *et al*., 2017). The dataset presented here uniquely contributes to this question by greatly improving taxon sampling of transcriptomes in a major plant clade (169 species in Caryophyllales) whose evolutionary history spans a time period that encompasses both deep divergences and more recent events during the Neogene (Smith *et al*., 2017). Likewise, our improved homology search and filtering approaches aid in identifying and pinpointing the phylogenetic locations of paleopolyploidy events. Moreover, we consider multiple lines of evidence for pinpointing the phylogenetic locations of paleopolyploidy events, including orthogroup tree topology, Ks plots, and chromosome counts. These approaches identified 26 paleopolyploidy events across Caryophyllales, include ten newly reported and 16 previously identified (Yang *et al*., 2015; Walker *et al*., 2017b). Importantly, two of these 26 events are suggested to be paleo-allopolyploidy events.

### Species trees based on transcriptome data are concordant with previous analyses

The species trees we recovered are highly concordant with previous analyses of family-level relationships (Cuénoud *et al*., 2002; Brockington *et al*., 2009; Schäferhoff *et al*., 2009; Arakaki *et al*., 2011; Yang *et al*., 2015). As seen in previous analyses, the placements of Sarcobataceae and Stegnospermataceae remain poorly supported despite using hundreds of loci of peptides or CDS sequences. We also discovered weaker support for two nodes that had previously received high support using a small number of loci. (1) Previous analyses recovered Cactaceae as being sister to Portulacaceae with 100% bootstrap support (Arakaki *et al*., 2011), but we recovered Anacampserotaceae+Portulacaceae as sisters to Cactaceae (modified phylome ICA = 0.31 and ASTRAL multi-locus bootstrap = 100). (2) Previous studies recovered strong to moderate support for the monophyly of Portulacineae+Molluginaceae [likelihood bootstrap = 100 (Arakaki *et al*., 2011), Bayesian posterior probability = 0.94 and parsimony bootstrap < 50 (Nyffeler & Eggli, 2010)], but we found low support for the relationship (ICA = 0.06 and multi-locus bootstrap = 93). This confirms that while individual loci can be informative, there is a large amount of phylogenetic conflict among gene trees (Smith *et al*., 2015; Walker *et al*., 2017b). Future studies should dissect these cases of discordance using a gene-by-gene approach (Brown & Thomson, 2016; Arcila *et al*., 2017; Shen *et al*., 2017; Walker *et al*., 2017a).

### Many paleopolyploidy events are associated with taxonomic units and/or habitat shifts

A notable pattern emerged showing that many paleopolyploidy events occurred on branches leading to major taxa and/or involved clear habitat shifts (Fig. 3). For example, the paleopolyploidy event within Nyctaginaceae is located on the branch representing a transition from trees and large shrubs in wetter environments within the Neotropics to a radiation of arid- and semiarid-adapted herbs and subshrubs recognized as Tribe Nyctagineae (Douglas & Manos, 2007; Douglas & Spellenberg, 2010). Similarly, the paleopolyploidy event at the base of Portulacineae is associated with the evolution of succulence (Nyffeler *et al*., 2008; Edwards & Ogburn, 2012; Ogburn & Edwards, 2013). Additional paleopolyploidy events are inferred along the branch leading to Polygonaceae (Schuster *et al*., 2013) and the branch leading to Droseraceae, a carnivorous family (Rivadavia *et al*., 2003).

Similar cases of paleopolyploidy events at or near the base of major clade origin include at the base of seed plants (Jiao *et al*., 2011), angiosperms (Jiao *et al*., 2011), monocots (Jiao *et al*., 2014), early eudicots (Jiao *et al*., 2012), and Asteraceae (Barker *et al*., 2016; Huang *et al*., 2016). This hints potential relationship between genome duplication and evolutionary innovations (Soltis & Soltis, 2016). On the other hand, however, branches leading to major recognizable taxonomic units tend to be relatively long and thus had more time to accumulate changes. Hence, while correlations between paleopolyploidy and evolutionary novelty are intriguing, we must be cautious in assuming that paleopolyploidy is the cause of such innovation (Smith *et al*., 2017).

### Inferring paleo-allopolyploidy from transcriptome data

Methods of paleopolyploidy detection developed for genomes or low-copy nuclear loci are inadequate for datasets with isoforms and missing duplicated copies (Lott *et al*., 2009; Jones *et al*., 2013; Marcussen *et al*., 2015; Thomas *et al*., 2016). While we applied a stringent filter to minimize missing taxa in orthogroups (no more than 2 missing), differential gene loss or silencing following polyploidy events remained a problem. Given that our goal of accurately pinpointing the phylogenetic location of paleopolyploidy events and that searching for allopolyploidy is highly dependent on taxon sampling, we only explored those cases when Ks vs. orthogroup tree-based mapping disagreed with each other.

Two allopolyploidy events were inferred in this study. We inferred the AMAR1 event (Fig. 2c) followed by a nested paleopolyploidy event (AMAR2) were together responsible for the observed Ks peaks, instead of a single, deeper event as previously reconstructed (Yang *et al*., 2015). *Schiedea* also has a complex history (Fig. 2d). While the polyploid origin of *Schiedea* was previously identified (Kapralov *et al*., 2009; Yang *et al*., 2015), by including its close relatives, *Honckenya* and *Scleranthus*, we show that all three species each had their own lineage-specific paleopolyploidy event, and that *Schiedea* likely had a yet unknown parental lineage other than the lineage leading to *Honckenya* (see schematic phylogram in Fig. 2d). The putative allopolyploid origin of *Schiedea* adds to a growing list of Hawaiian endemic radiation with similar putative allopolyploid origins (Barrier et al., 1999; Yang & Berry, 2011; Marcussen et al., 2012; Roy et al., 2015), and demonstrates the importance of information concomitant with increased transcriptomic taxon sampling. Moving forward genome and transcriptome data will be essential for investigating selection, homeolog expression, gene silencing and loss in contributing to these divergence events following allopolyploidy.

### Improved homology inference methods improve paleopolyploidy mapping

In the original phylome approach (Huerta-Cepas *et al*., 2011), each sequence from a seed species was used to search against a database of sequenced genomes. The resulting homologous sequences were filtered, aligned, and phylogenetic trees were constructed. In this study, we made three modifications that enhance the ability to accommodate transcriptome data. First, by merging putative homolog groups that represent gene duplication within Caryophyllales, we ensure that each of our final orthogroup contains a distinctive lineage that diversified from a single copy at the MRCA of Caryophyllales. Secondly, given the presence of multiple transcript isoforms and the inherent incompleteness of transcriptome datasets, we added a second step using transcriptome sequences as queries to search against the beet proteome for sorting transcriptome-derived sequences into backbone orthogroups constructed with genomes only. Lastly, to clean up spurious tips and isoforms, we added tip-trimming and long-branch cutting steps. By taking this two-step, baited approach we were able to process a large number of taxa without going through the time consuming all-by-all homology search required by OrthoMCL (Li *et al*., 2003) and OrthoFinder (Emms & Kelly, 2015). A second advantage of modified phylome is that it avoids a graph-based clustering step, and hence is not biased by sequence length or phylogenetic relatedness among taxa (Emms & Kelly, 2015). The modified phylome approach is more effective than other baited approach such as HaMStR (Ebersberger *et al*., 2009) in that it explicitly takes gene tree topology into account in distinguish ortholog from paralogs. However, because both the original phylome and the modified methodology start with a seed genome, the resulting orthogroup set is dependent on the quality and gene content of the focal genome.

Despite the efficiency of the modified phylome approach, phylogenetic uncertainty grows significantly with increased dataset size. To overcome this challenge, we took a divide-and-conquer approach and used a second homology inference strategy and inferred subclade species trees using all-by-all homology search, Markov Clustering (van Dongen, 2000) of filtered hits, followed by alignment and tree trimming (Yang & Smith, 2014). We use the original Markov Clustering (MCL) instead of software packages like OrthoMCL (Li *et al*., 2003) and OrthoFinder (Emms & Kelly, 2015) that aim directly at obtaining orthogroups using filtered and normalized BLAST hits. The normalization procedures used by these software packages were based on genome-derived data, and were not evaluated using transcriptome datasets that had isoforms, missing data, and assembly errors. By using the original MCL with relatively low inflation value (i.e. coarse clusters) and taking advantage of outgroup information to root and extract orthogroups, we are able to minimize the loss of gene duplication information in our dataset.

Our techniques inferred consistent percentages of gene duplication as compared to analyses using plant groups of similar ages (Barker *et al*., 2016; Huang *et al*., 2016; Xiang *et al*., 2016). We found that within inferred paleopolyploidy events, approximately one-third of genes display clear evidence of duplication (i.e., they retain at least two overlapping taxa between paralogs), similar to the numbers we found previously in both transcriptomes and genomes (Yang *et al*., 2015). For example, the PHYT2, AMAR1, and AMAR3 events follow this “one-third rule” (Fig. 2). When there is phylogenetic uncertainty, gene duplication events may be mapped to two adjacent nodes, each with lower percentages, such as observed for PORT1 and CARY1. Percentages of gene duplication can be inflated during rapid diversifications, where short internodes and phylogenetic uncertainty make it difficult to distinguish paralogous copies from isoforms using tree topology. Such inflated percentages of gene duplication can be seen at the base of Cactaceae and *Silene* (without paleopolyploidy), and at PHYT1 and NCORE1 (following a paleopolyploidy event).

Moving forward, additional taxon sampling of genomes and transcriptomes will be essential to identify additional paleopolyploidy events and pinpoint their phylogenetic locations. Understanding the legacy of ancient polyploidy events in plant macroevolution will require many other forms of improved data as well, including functional studies of traits, molecular pathways, and genes themselves. Only then will we have a more comprehensive functional framework for understanding differential gene retention and diploidization following paleopolyploidy events, and how paleopolyploidy is linked to character evolution and niche shifts.

## Acknowledgements

The authors thank H. Flores Olvera, H. Ochoterena, N. Douglas, S. Lavergne, T. Stoughton, L. Crawford, G. Kadereit, R. Puente, L. Majure, M. Howard, D. Anderson, M. Palmer, and K. Thiele for assisting with obtaining plant materials; the Cambridge University Botanic Gardens, Bureau of Land Management, White Sands Missile Range, U.S. Forest Service, California State Parks, Rancho Santa Ana Botanic Garden, Desert Botanical Garden, Millennium Seed Bank, and Booderee National Park for granting access to their plant materials; and M. R. M. Marchán-Rivadeneira, and M. Parks for help with laboratory work. The molecular work of this study was conducted in part in the Genomic Diversity Laboratory at the University of Michigan. Support came from the University of Michigan, Oberlin College, the National Geographic Society, the U.S. National Science Foundation (DEB 1054539, DEB 1352907, and DEB 1354048), and a Natural Environment Research Council Independent Research Fellowship (NERC RG69516).

## Author contribution

S.A.S., S.F.B., M.J.M., and Y.Y designed research; M.J.M., Y.Y., J.M., J.M., J.O., S.F.B., and J.F.W. collected samples and carried out lab work; Y.Y. analyzed data and led the writing. All authors edited the manuscript and approved the final version.

## Supporting Information

**Fig. S1.**
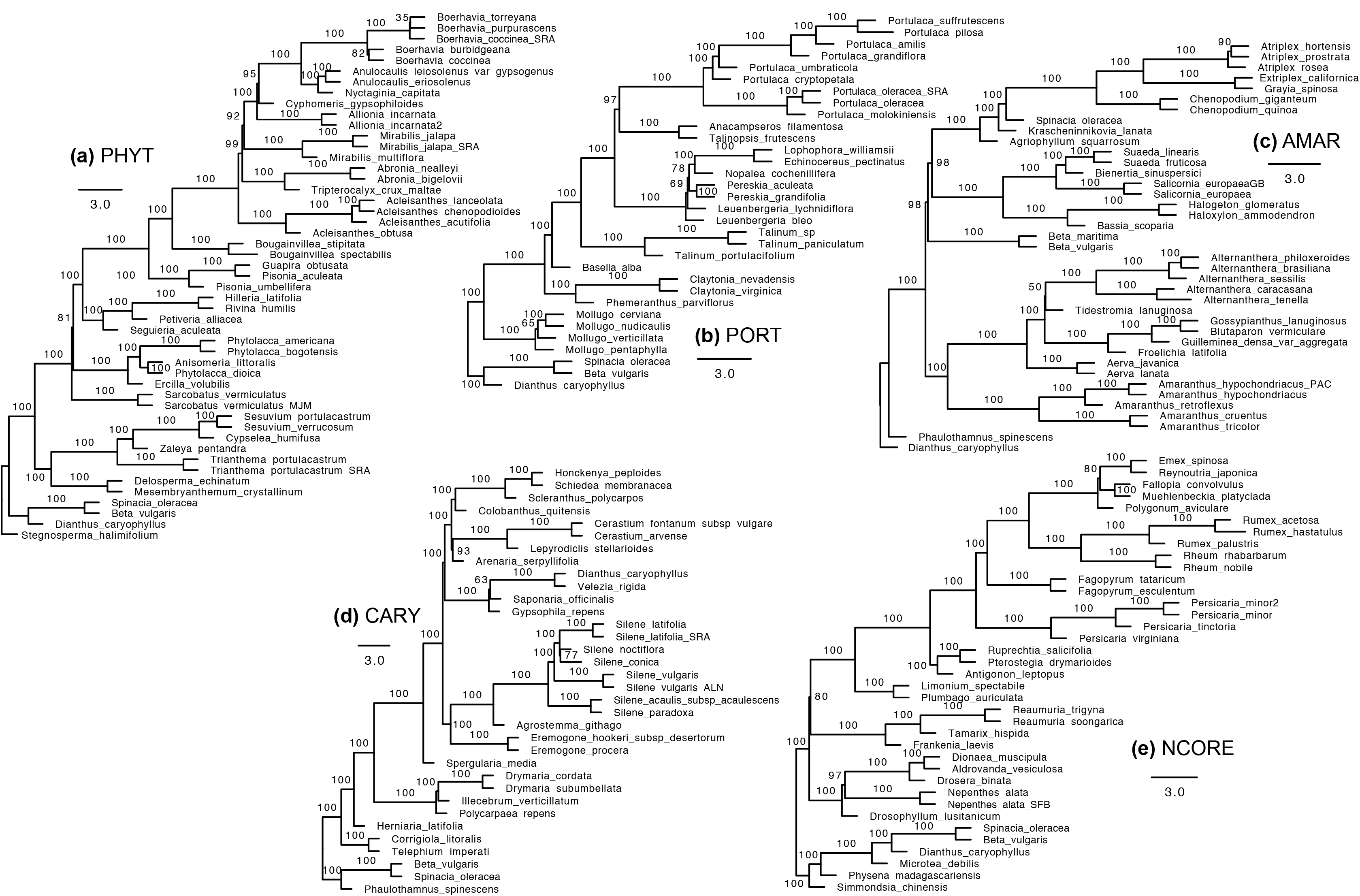
Subclade Maximum Quartet Support Species Tree (MQSST) from ASTRAL analysis of individual subclade orthologous gene trees. Numbers on branches are multi-locus bootstrap percentages. Branch length represents coalescent units, with terminal branch lengths artificially fixed to 1 as coalescent units cannot be calculated for terminal branches.

**Fig. S2.**
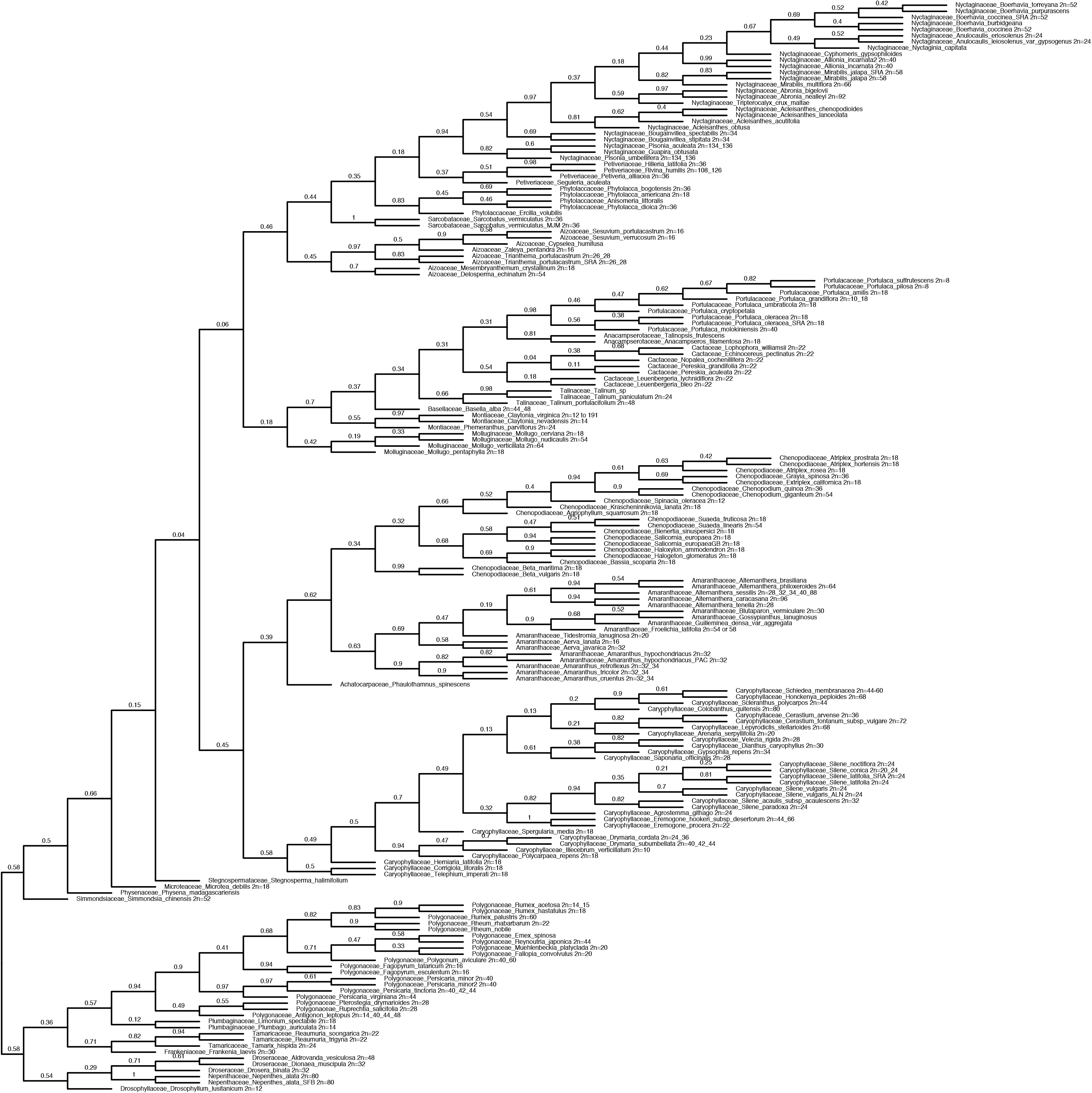
Phylogram from RAxML analysis of the concatenated 624-gene supermatrix from genome walking, with ICA scores on branches.

**Fig. S3.**
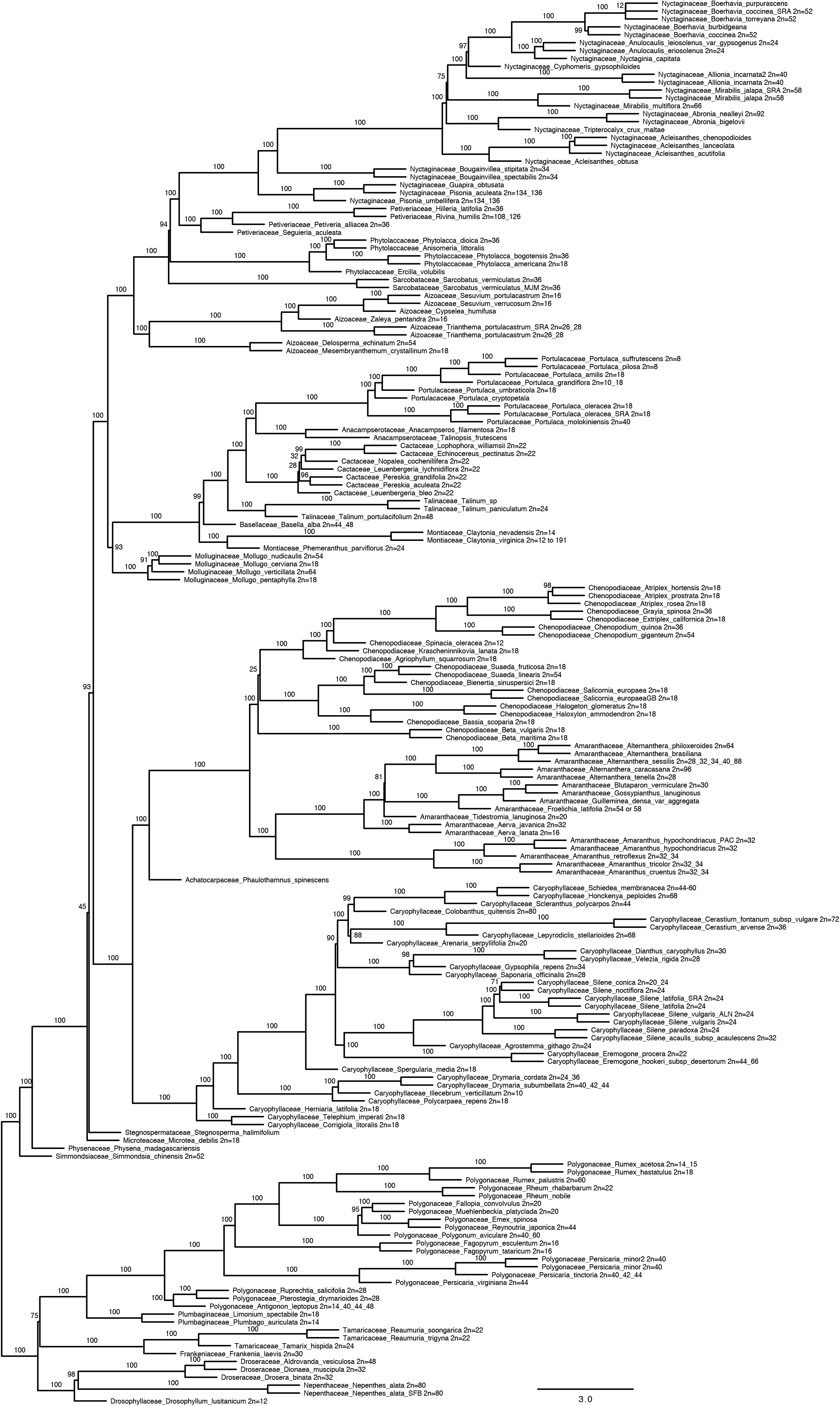
Subclade Maximum Quartet Support Species Tree (MQSST) from ASTRAL analysis of 624 orthologous gene trees from genome walking. Numbers on branches are multi-locus bootstrap percentages. Branch length represents coalescent units, with terminal branch lengths artificially fixed to 1 as coalescent units cannot be calculated for terminal branches.

**Fig. S4.**
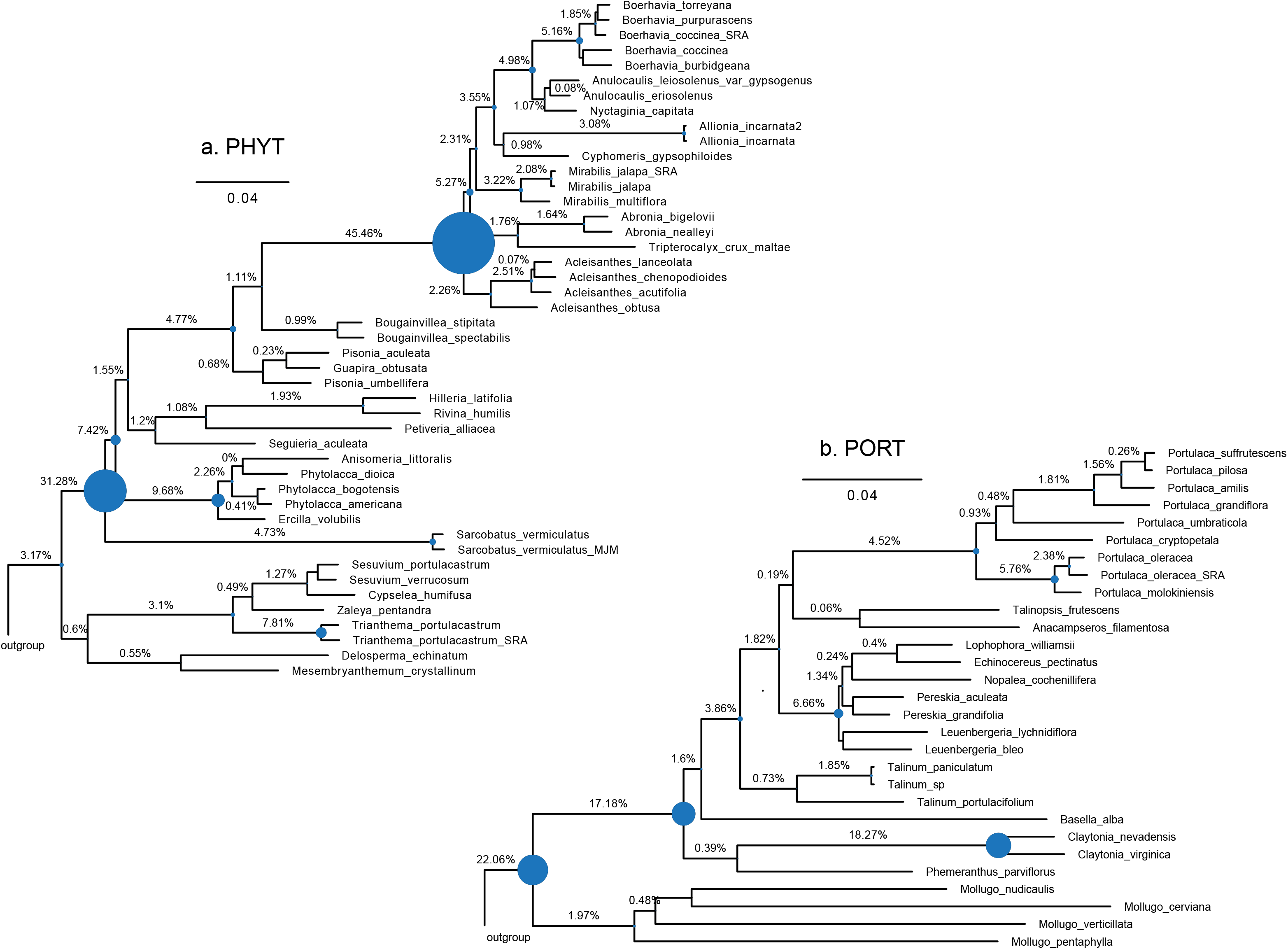
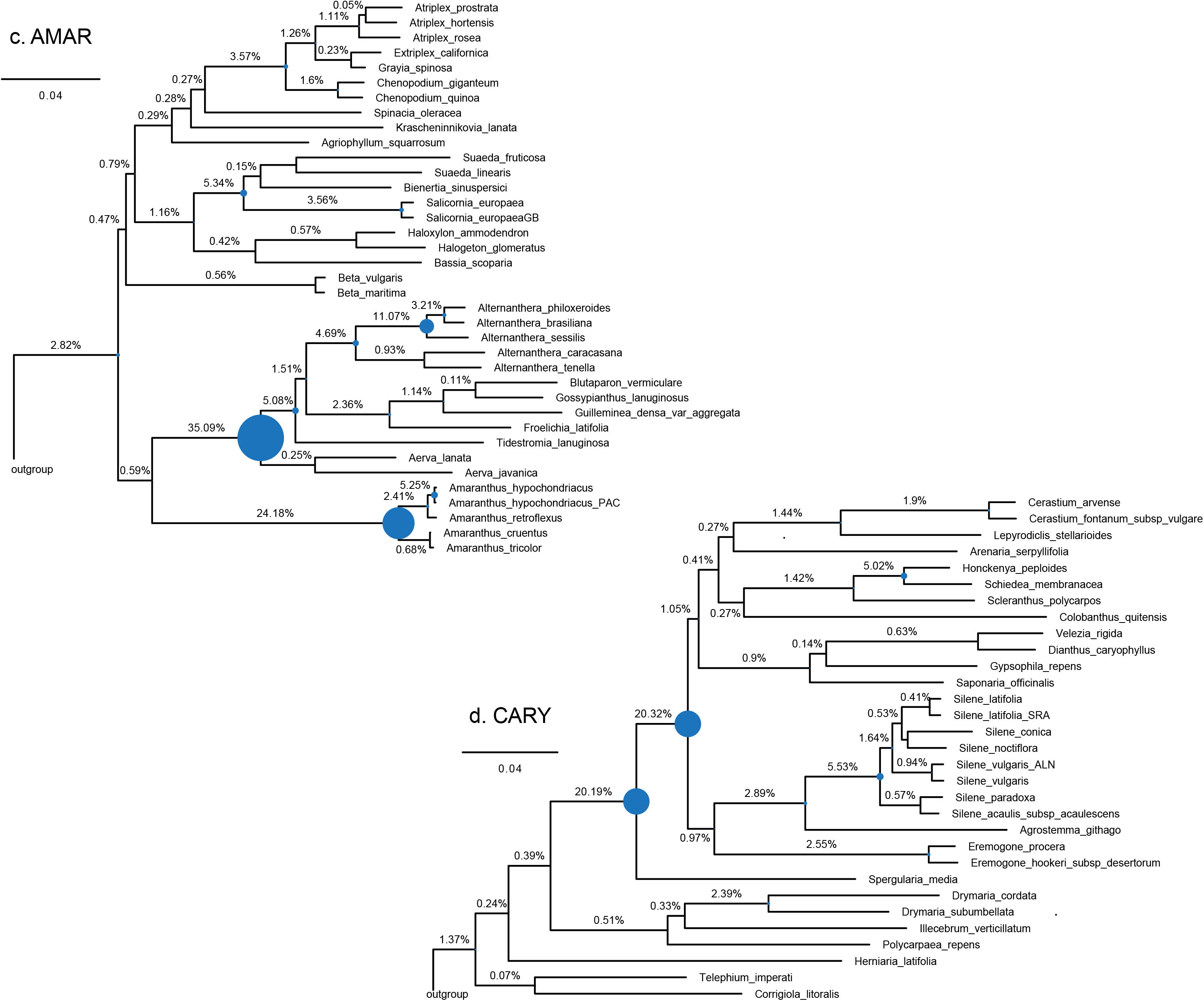
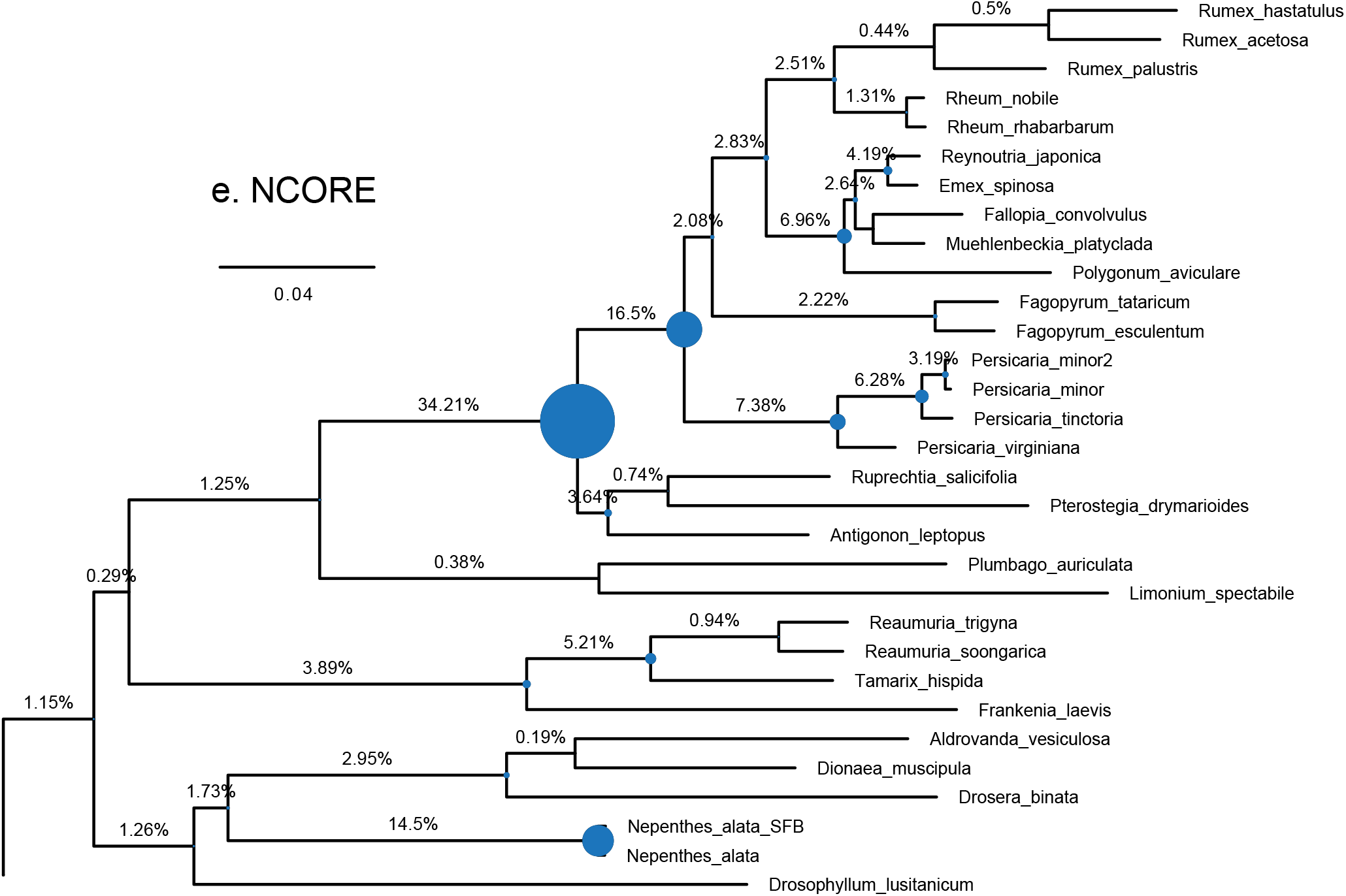
Proportion of orthogroups showing duplications, filtered by local tree topology in addition to orthogroup-wide average bootstrap support. Bubble diameters are proportional to frequency of gene duplication.

**Fig. S5.**
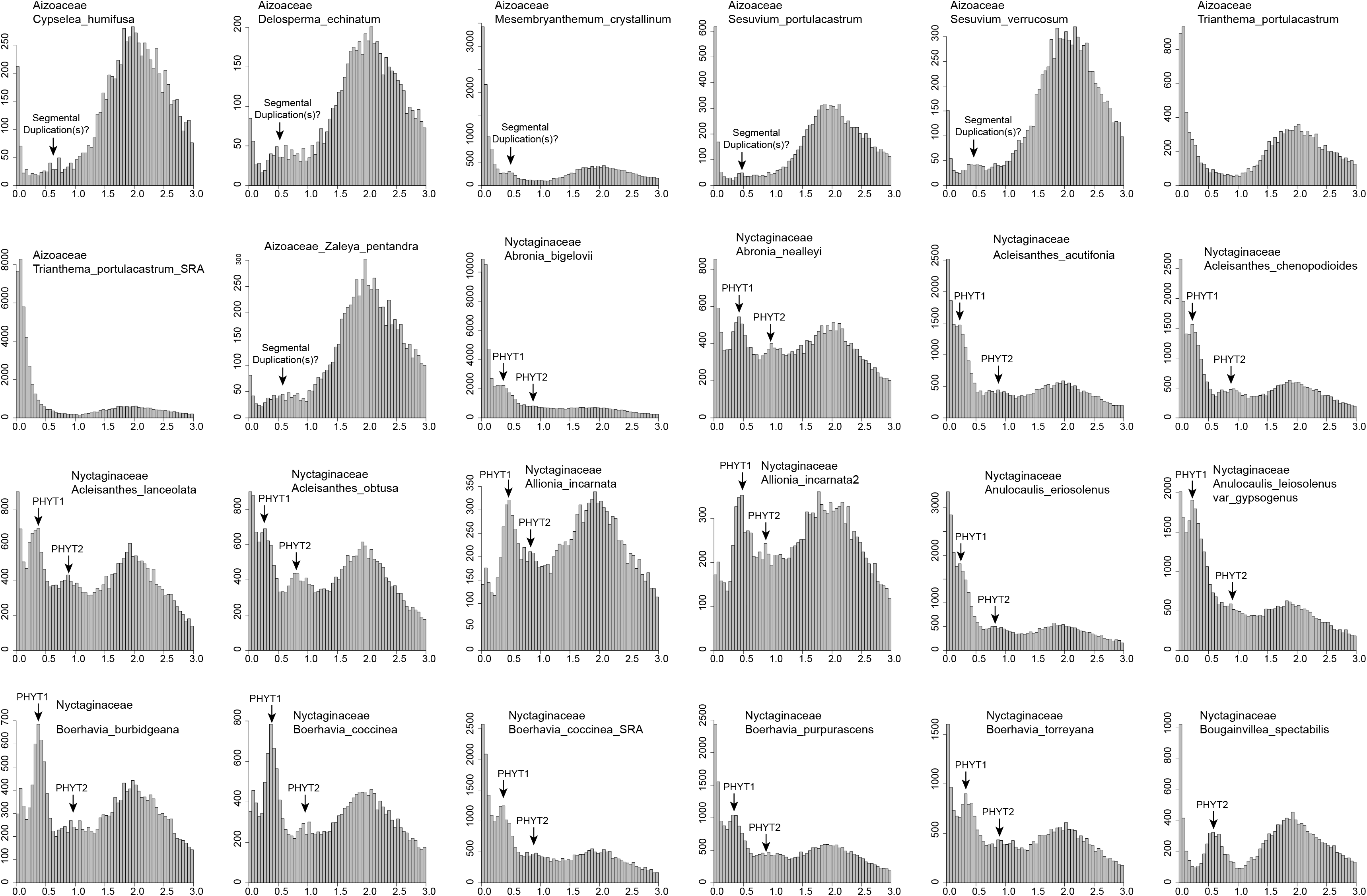
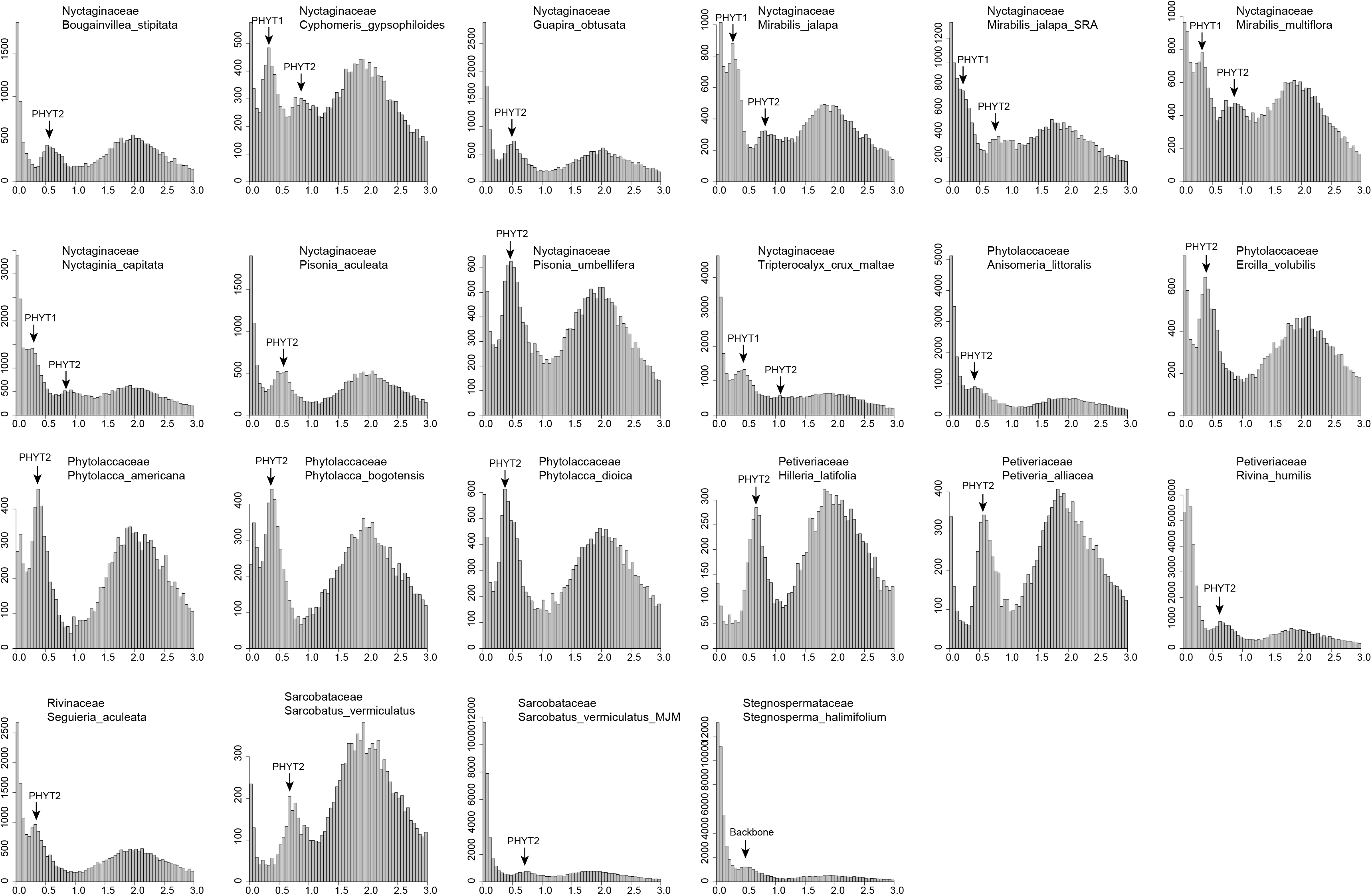
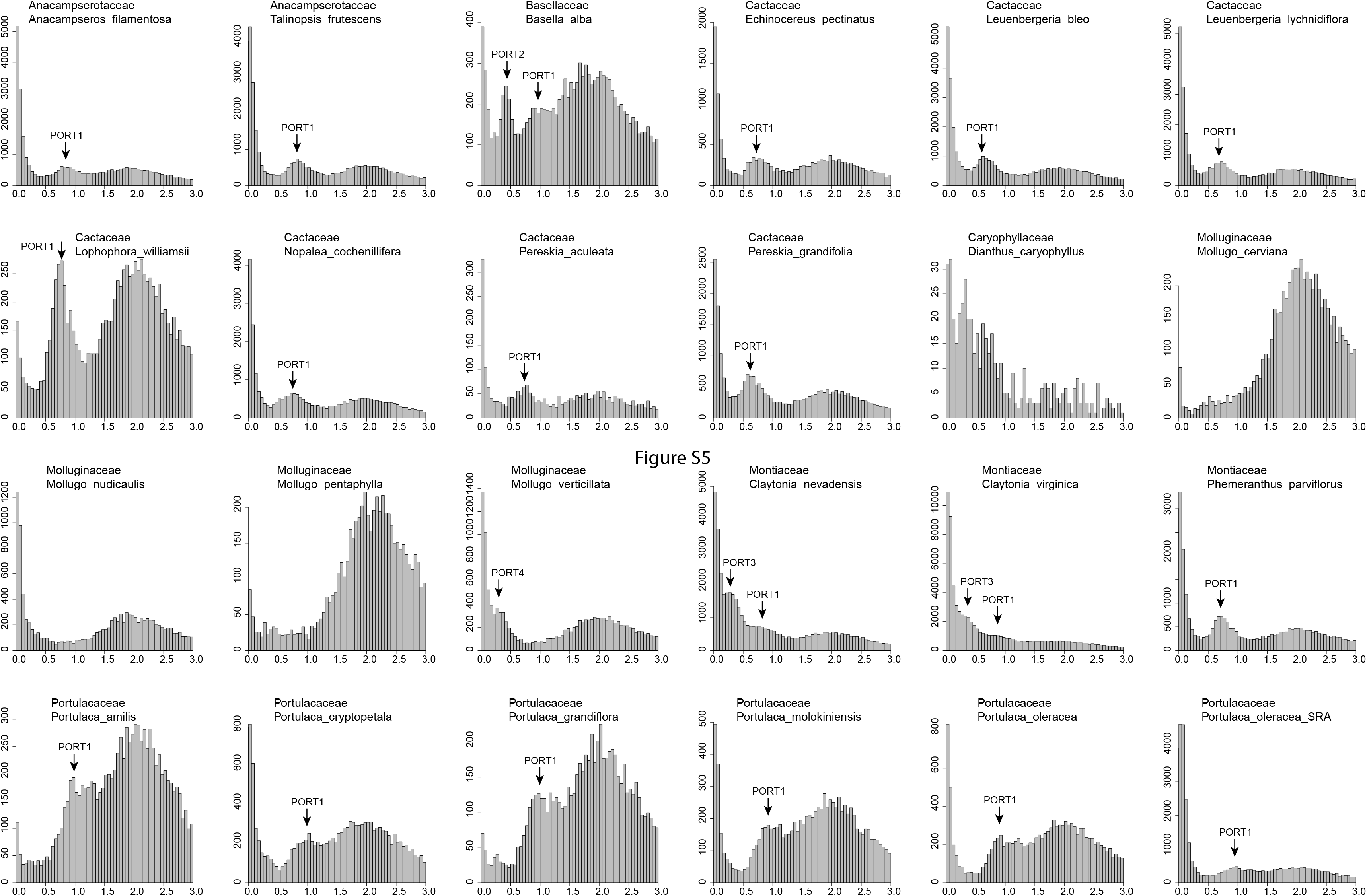
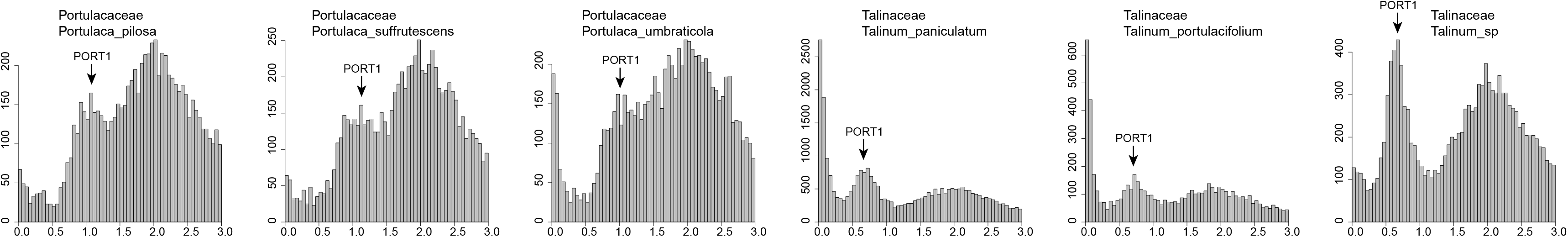
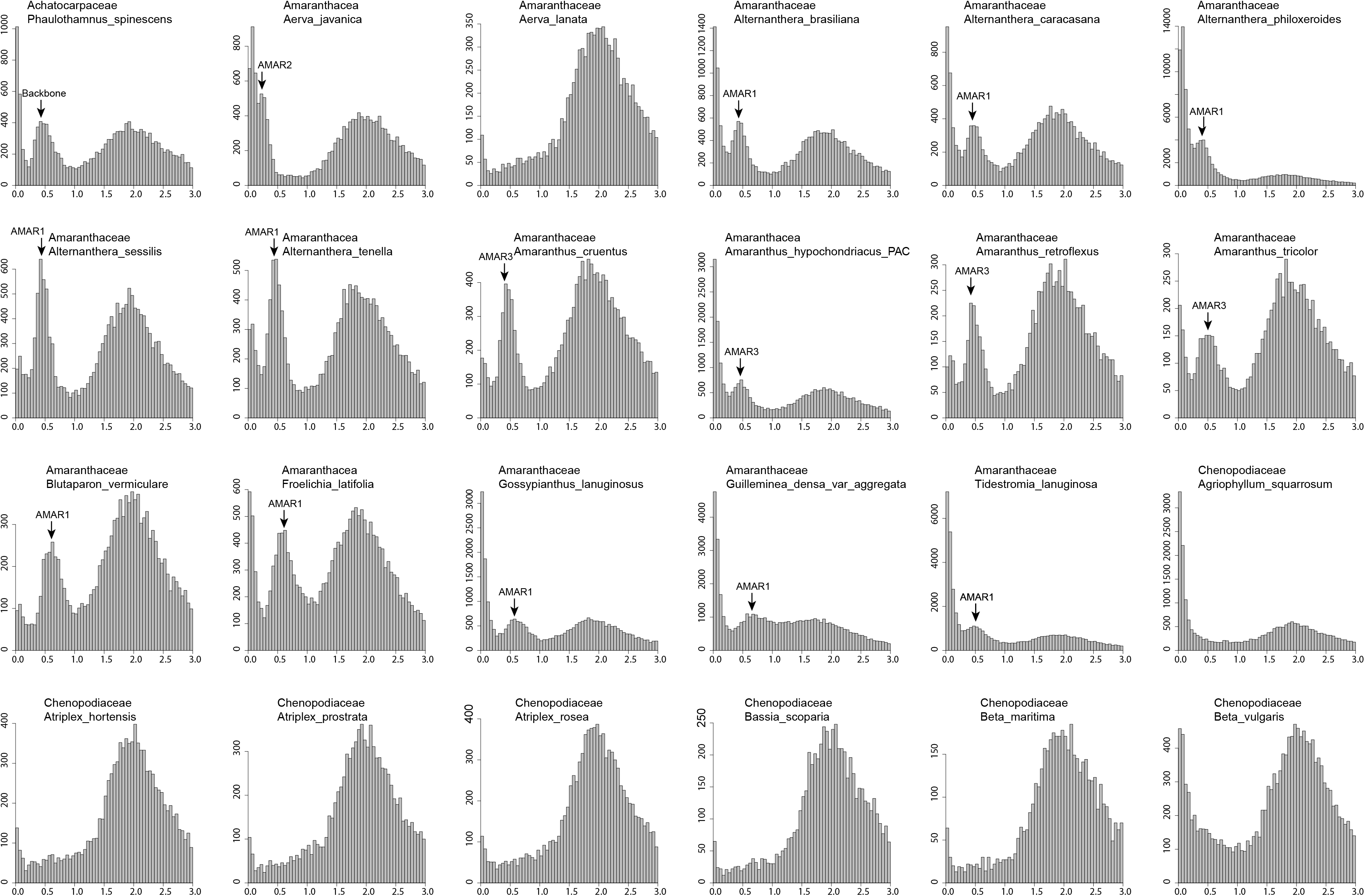
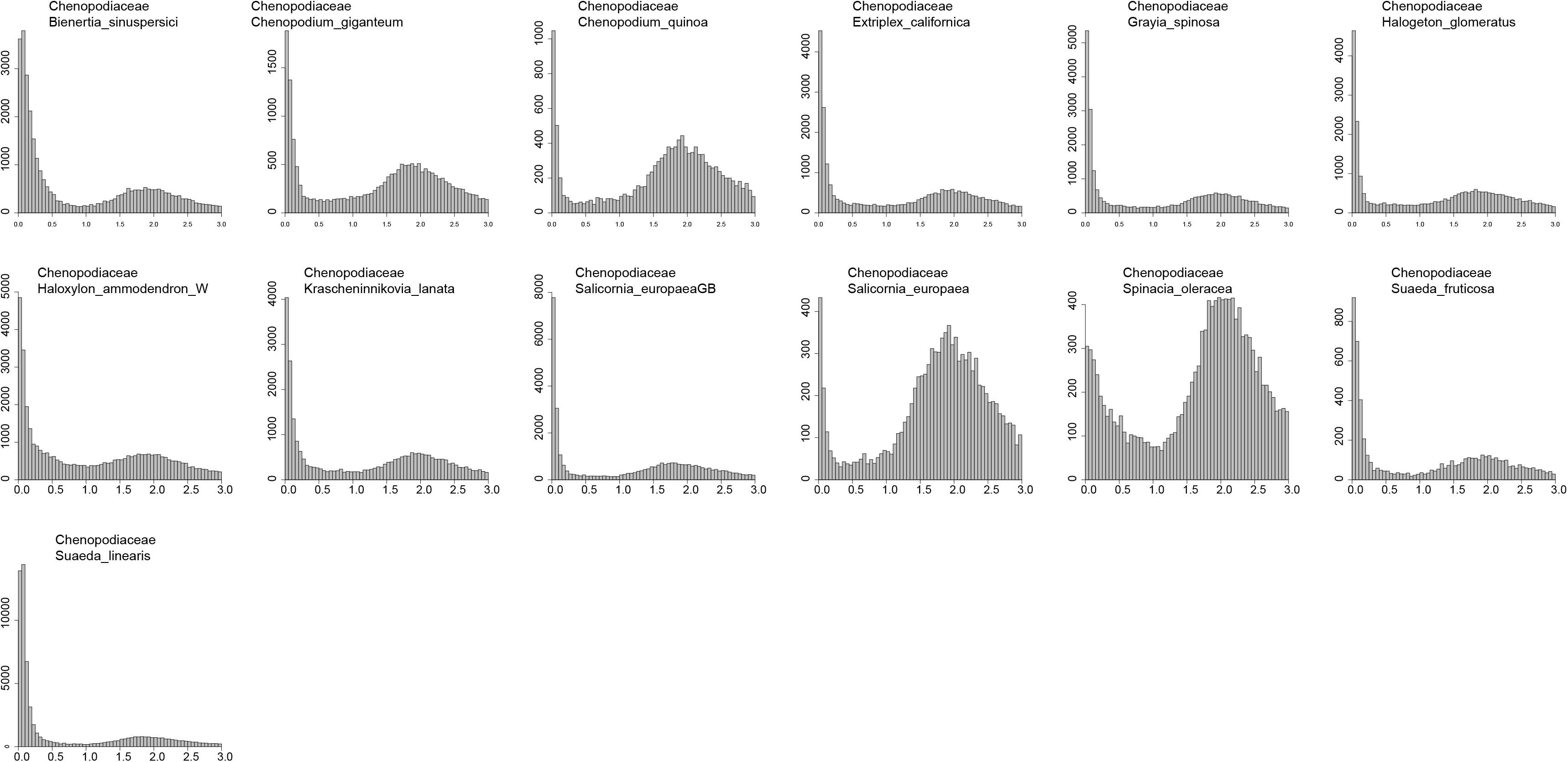
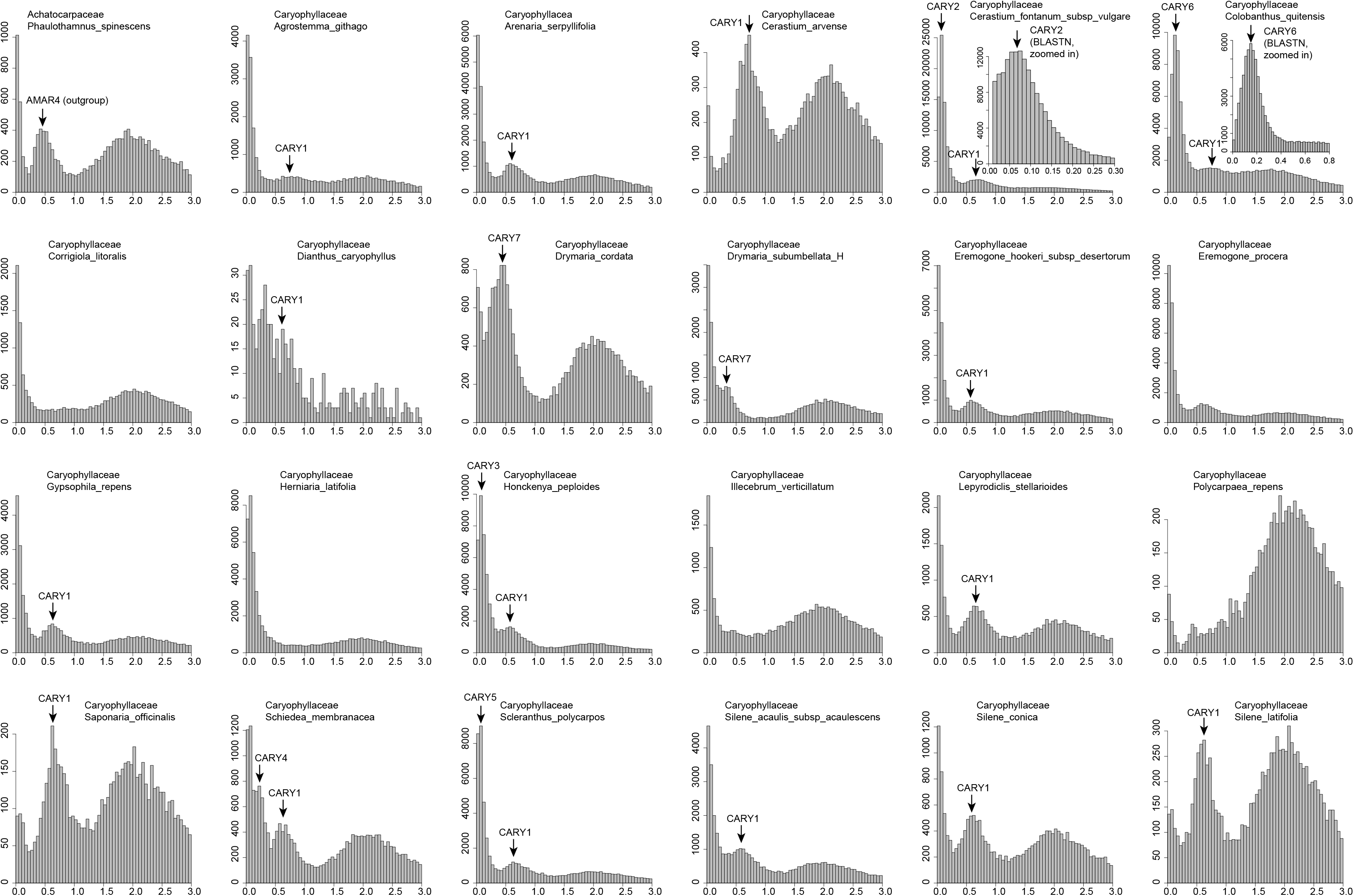
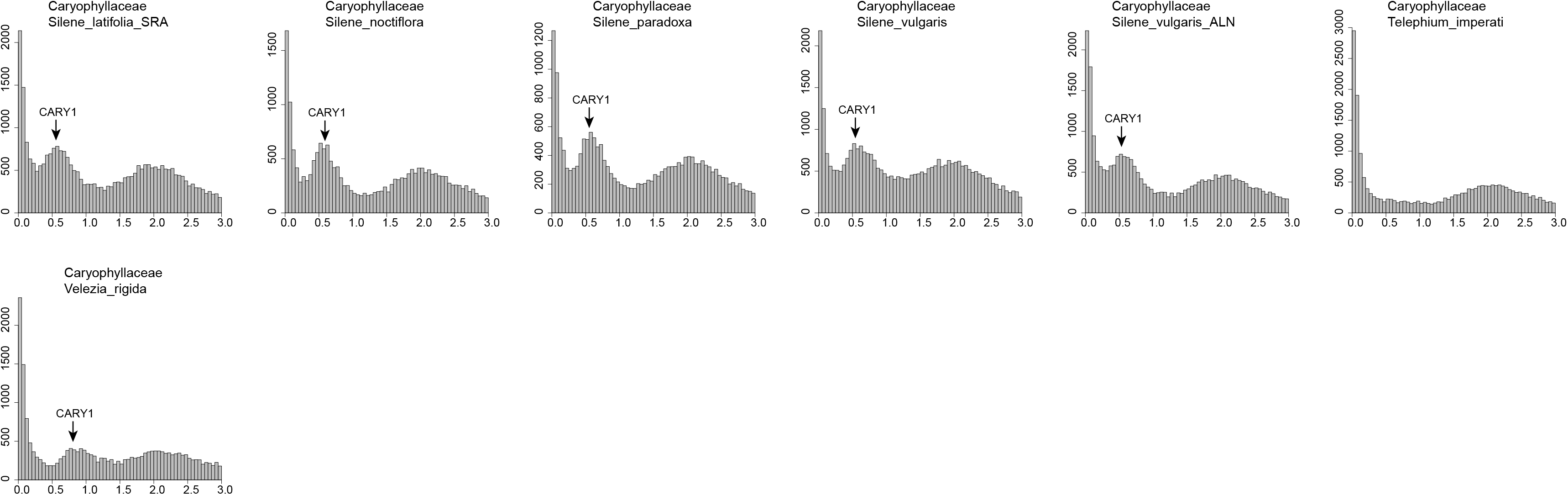
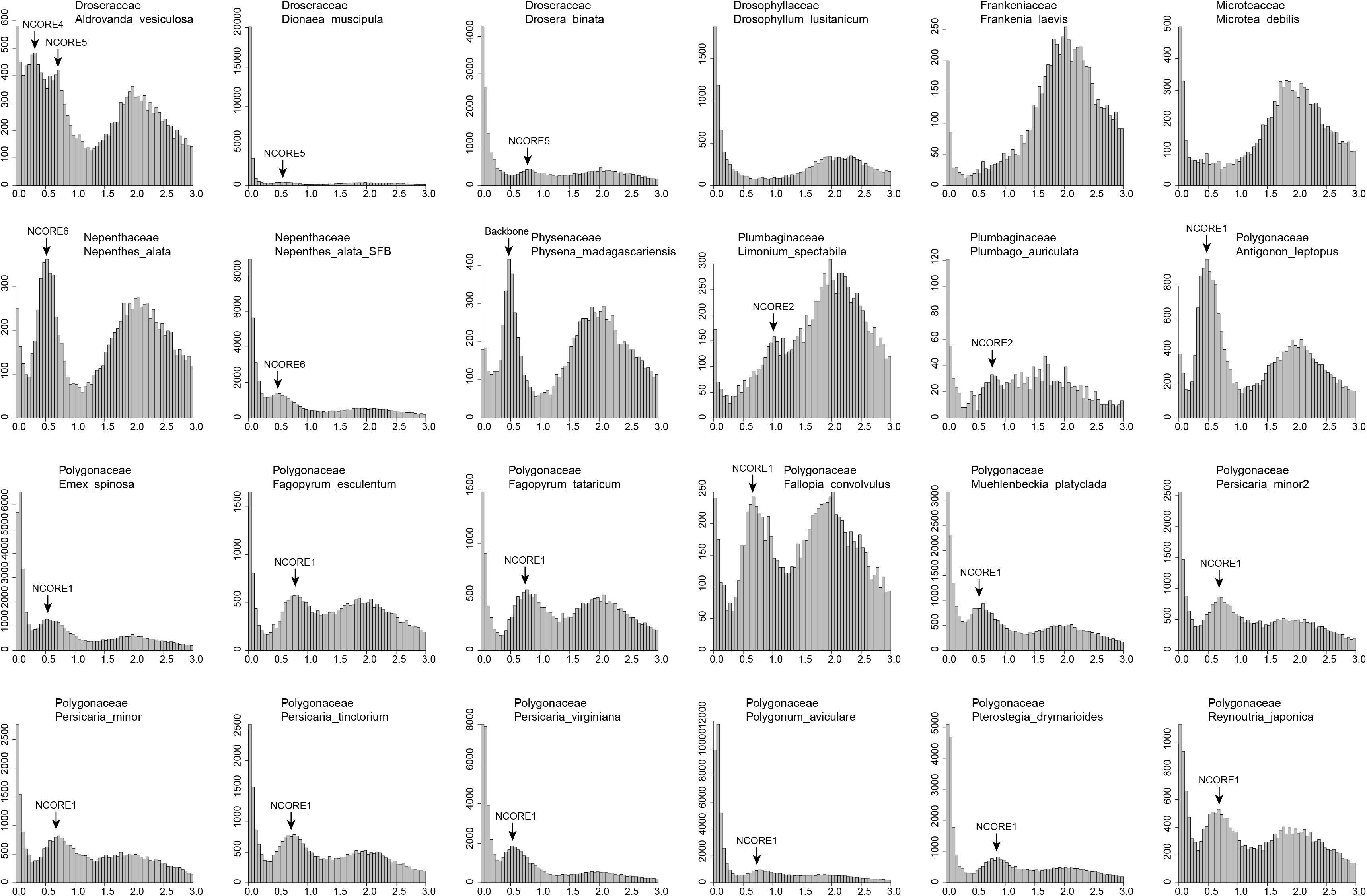
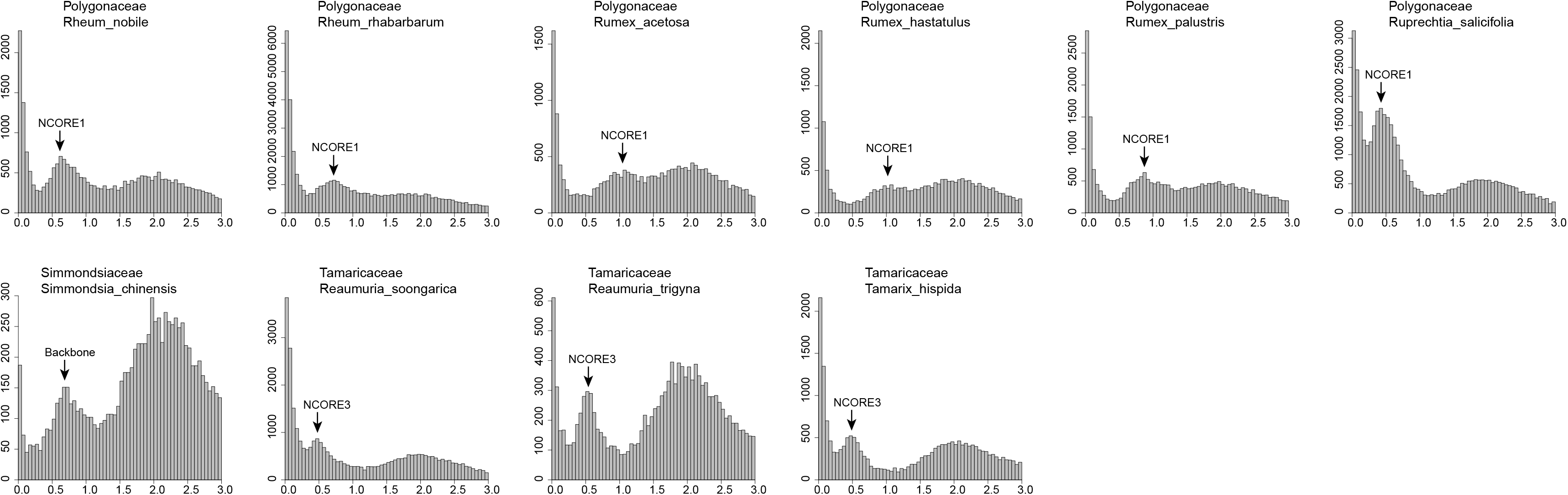
Distribution of synonymous substitutions (Ks) for Caryophyllales paralogs based on BLASTP hits. Zoomed-in Ks plots from BLASTN hits are shown in addition to the results from BLASTP hits to show more recent Ks peaks.

**Table S1** Sources of data and settings for read cleaning, assembly, and translation.

**Table S2** Information for 51 newly sequenced transcriptomes Methods S1 All-by-all homology search for subclade datasets. Information for the 43 newly sequenced transcriptomes. YY=Ya Yang, MJM=Michael J. Moore, SFB=Samuel F. Brockington, JM=Jessica Mikenas, JO=Julia Olivieri, JFW=Joseph F. Walker. Details for collection, RNA extraction, and library preparation protocols can be found in Yang et al. (2017)

**Methods S1** Coding sequences from each transcriptome were reduced using cd-hit-est (-c 0.99-n 10) as part of the CD-HIT package. Homology searches were carried out using all-by-all BLASTN with an E value cutoff of 10 and max_target_seqs set to 1000. BLASTN output was filtered by requiring the alignment to cover 40% of both the query and hit sequences. Clustering was performed using Markov CLuster algorithm [MCL v12-068 (van Dongen, 2000; van Dongen & Abreu-Goodger, 2012)] from filtered hits with the E value cutoff set to 10^−5^ and an inflation value of 1.4. Ends of sequences with no interspecific BLASTN hit coverage were clipped. Remaining sequences shorter than 40 characters were removed, and clusters with more than 2 taxa missing were ignored. Each resulting cluster was aligned, alignments were trimmed as in genome walking, and phylogenetic trees were estimated with RAxML using the model GTRCAT. Spurious tips were trimmed using a relative tip cutoff of 0.2 and an absolute tip length cutoff of 0.4 (0.3 and 0.6 for NCORE and 0.1 and 0.3 for CARY, respectively), and isoforms were removed with the same criteria as genome walking. Long internal branches longer than 0.3 (0.4 for NCORE and 0.2 for CARY and AMAR) were separated. The same procedure was repeated once before producing the final homolog trees with 200 fast bootstrap replicates.

